# From mice to humans: A multi-omic predictive framework for translational immunology

**DOI:** 10.64898/2026.09.18.752716

**Authors:** Wasim Aluísio Prates-Syed, Aline A Lira, Nelson Cortes, Jaqueline DQ Silva, Bárbara Hamaguchi, Evelyn Carvalho, Adriana Castillo-Chávez, Ricardo Durães-Carvalho, Otavio Cabral-Marques, Ester Cerdeira Sabino, José Eduardo Krieger, Thomas Hagan, Gustavo Cabral-Miranda

**Author notes:** **Correspondence to:** or, Gustavo Cabral-Miranda, Laboratory of Genetics and Molecular Cardiology, Heart Institute, Clinical Hospital, Faculty of Medicine, University of São Paulo, Brazil. And Institute of Tropical Medicine, Faculty of Medicine of the University of São Paulo, Brazil. Equal contribution as senior authors (Thomas Hagan, Gustavo Cabral-Miranda).

## Abstract

Mice are key preclinical animal models in vaccine and immunological research, yet their predictive value for human immunity remains contested. Here, we evaluated the translatability of murine models across inactivated and subunit vaccination (influenza, hepatitis B), acute infection (*S. aureus*, *E. coli*), and injury (burns and trauma), integrating transcriptomic profiles with sequence evolution, cis-regulatory architecture, and functional annotation. Functional modules were more conserved between species than individual orthologous genes. Translational accuracy depended on stimulus intensity, as infections and injuries engaged conserved signatures, while single-dose vaccination diverged. We then built multilayer models to predict human expression rank and direction of change, and to classify shared leading-edge genes. Adding evolutionary and regulatory layers improved these predictions. We provide a step-by-step R Markdown notebook to apply the models to user data. The study code and datasets are available at https://github.com/wapsyed/mousetohuman_multilayer

**GRAPHICAL ABSTRACT:** 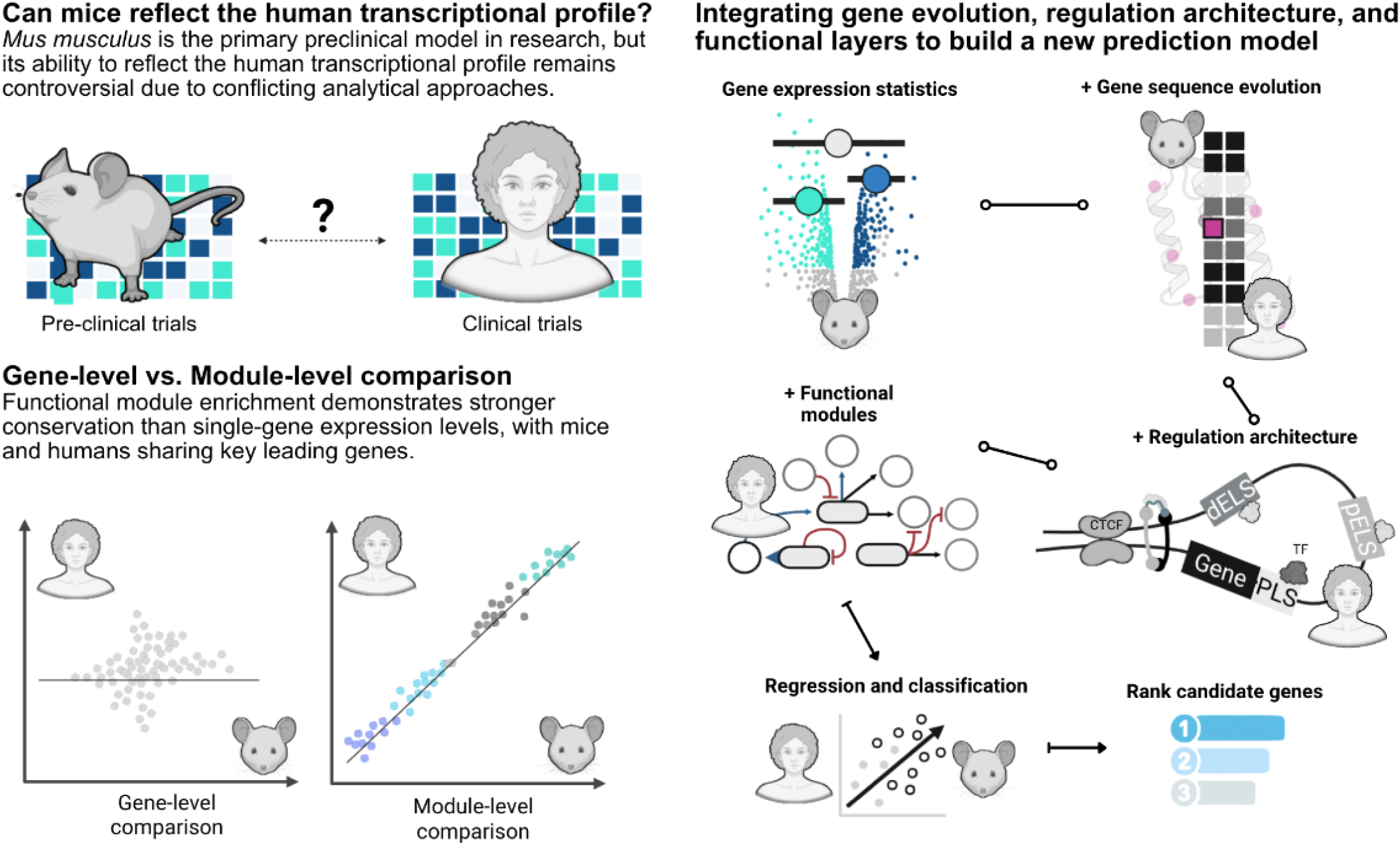

## 1. INTRODUCTION

Vaccine development over the past two centuries represents one of the most significant achievements in modern medicine, providing direct protection through immune responses ^1^. However, the growing complexity of the immune system underscores the need for continued research to fully understand its mechanisms. While vaccines are designed to trigger specific responses, a key challenge remains to determine how these responses differ across species, particularly between humans and mice, the most widely used animal model in preclinical studies^2^.

Mice are an essential tool for biomedical and immunological research, valued for their genetic tractability, short lifespans, and physiological similarities to humans ^3,4^. Mice and humans share most of the recently evolved adaptive immune system and exhibit high conservation in protein-coding regions, alongside many other genomic and phenotypic similarities, including cytometry-defined cell populations ^5,6,7,8,9,10^.

However, the biology of these species can widen the gap between preclinical and clinical research, contributing to clinical trial failures and raising questions about the predictive value on vaccination ^11,12,13,14^. These differences include low conservation in non-protein-coding sequences ^15^ and in homologous tissues ^16^, among others, as reviewed elsewhere ^7^. Furthermore, the temporal dynamics of immune responses are often overlooked in cross-species comparisons. The assumption that a given time point in a murine model corresponds linearly to the same chronological time in humans ignores potential physiological differences in response kinetics.

Given these differences, the translational relevance of mouse models has been a subject of considerable debate ^14,17,18,19,20^. However, these studies did not investigate why specific genes maintain higher expression conservation than others, particularly under perturbation, and left statistical issues unaddressed, focusing on descriptive pathway overlaps without systematically connecting cross-species expression conservation to the underlying genomic and epigenomic architecture. Nonetheless, others have shown that regulatory architecture and functional transcription factor (TF) networks exhibit substantial conservation between homologous tissues across species, even when individual sequence-level transcription factor binding occupancies diverge ^16,7,10^.

Computationally, comprehensive databases such as ARCHS4, and prediction models such as Found in Translation (FIT) and semi-supervised neural-network-based tools were developed to bridge the gap between mice and humans and help characterize and predict human disease and immune responses from mice data, among other applications ^21,22^. In parallel, deep learning foundation models like GeneRAIN have leveraged hundreds of thousands of bulk transcriptomes to capture cross-species gene conservation beyond DNA sequence similarity ^23^. However, existing computational tools rely predominantly on single-gene linear models or unweighted aggregations that fail to capture the non-linear, multi-omic constraints, such as sequence evolution rates, TF binding, and cis-regulatory architecture, governing cross-species immune response translatability. Practically, their implementation often presents substantial accessibility and reproducibility barriers, as several key frameworks depend on proprietary platforms such as MATLAB or that lack fully open, customizable end-to-end retraining pipelines.

Leveraging the development of omics technologies, the field of systems vaccinology has deepened our understanding of human immune responses to vaccination. Our groups have recently developed vaccine atlases describing human transcriptional responses across multiple vaccines ^24,25^. Despite these resources, little work has been done to understand the overlap in transcriptional responses to vaccinations between mice and humans.

In this study, we systematically compare three stimulus categories, including inactivated and subunit vaccination (Fluad, Engerix B), acute infection (S. aureus, E. coli), and injury (burn, trauma-haemorrhage), to quantify mouse-to-human translatability and reconcile contradictory prior reports. We show that functional modules correlate more strongly than individual gene effect sizes or ranks, and that leading-edge genes (LEGs) ranking within conserved modules preserves cross-species rank information. We then integrate gene sequence evolution, cis-regulatory architecture, and machine learning to dissect why gene-level conservation fails despite module-level preservation, and why certain genes converge in their log2FC and ranking, offering a practical prioritisation framework for preclinical target selection. Rather than reiterating the well-established conservation of pathways over individual genes, this work introduces a quantitative framework that links stimulus intensity, analytical granularity, and multi-omic layers as interdependent determinants of translatability.

## 2. RESULTS

### 2.1. Data curation

From 201 mouse immunization datasets retrieved from GEO (**Fig. 1a**), only two microarray datasets matched to human vaccination data in our atlases, by tissue, antigen, vaccine type, target condition, time points, and adjuvants **(Supplementary Table 1, Fig. 1b)**. One dataset (GSE120661) included influenza and hepatitis B vaccinations, while the other (GSE182858) involved influenza vaccination in dirty mice ^26^, which was not used in this analysis. Therefore, only GSE120661 was carried forward. The dirty-mouse cohort could not be directly compared with the human cohort owing to divergent sampling schedules (3 hours before and after immunisation vs. days 1, 3, and 7, respectively) ^12^.

**Fig. 1.**
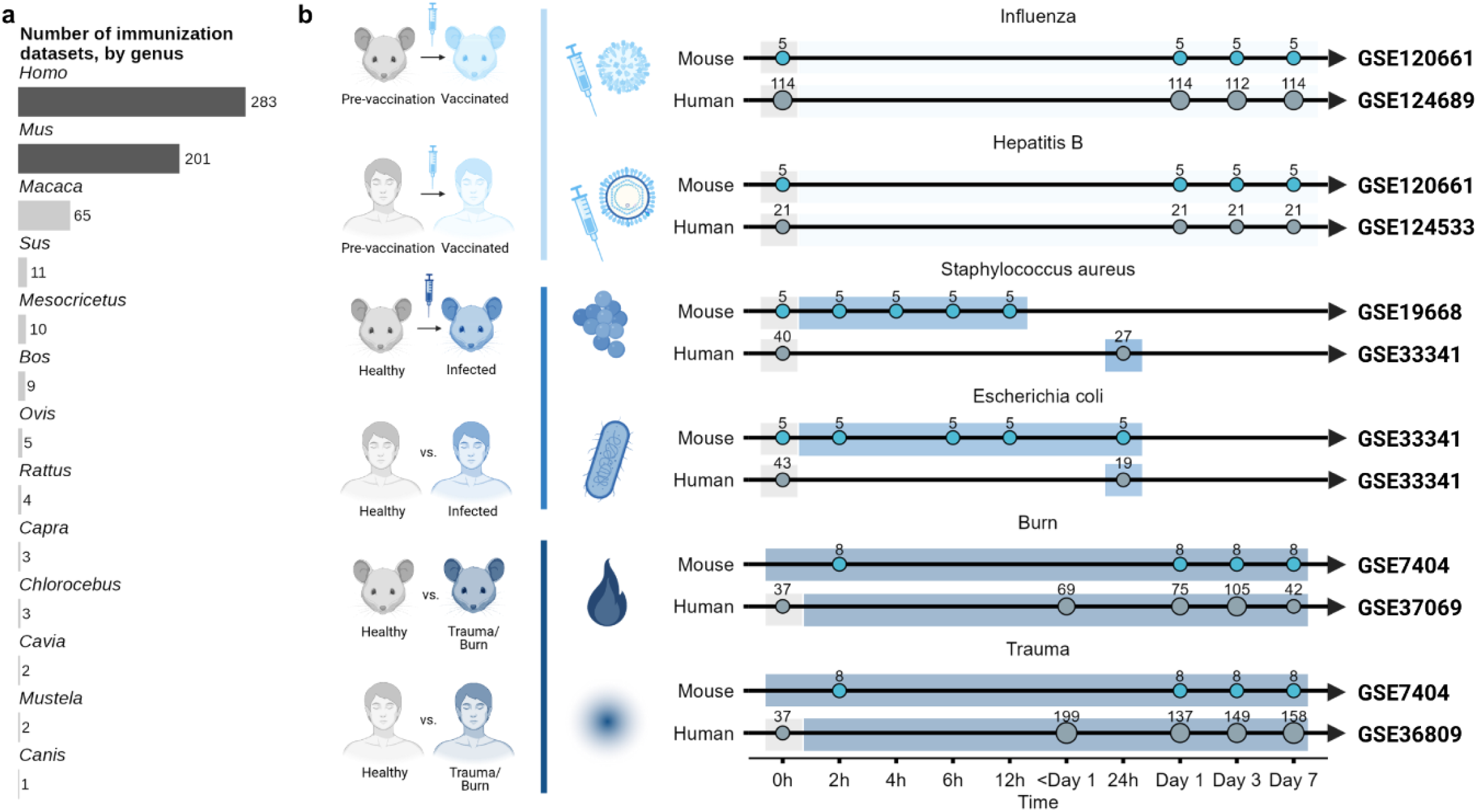
Study Overview and Data Curation. (a) Number of immunization datasets available by genus, showing the predominance of Homo and Mus (blue). (b) Experimental timelines for matched mouse and human datasets across the four conditions, namely influenza and hepatitis B vaccination, S. aureus and E. coli infection, and burn and trauma injury. Colours indicate control (grey), vaccinated (light blue), infected (blue) and injured (dark blue).

The GSE120661 dataset, comprising peripheral blood mononuclear cell (PBMC) samples collected up to 168 hours post-vaccination with the MF59-adjuvanted influenza vaccine Fluad®, was selected to evaluate the mouse model’s performance. This dataset closely matched the design of the human Fluad® dataset (GSE124689), which included PBMC samples collected at corresponding time points (days 1, 3, and 7). To broaden the scope of our analysis beyond vaccination, we also extended our curation to *Staphylococcus aureus* (GSE19668 for mouse, GSE33341 for human) and *Escherichia coli* infection datasets (GSE33341 for mouse and human), as well as injury conditions, including burns (GSE7404 for mouse, GSE37069 for human) and trauma-hemorrhage (GSE7404 for mouse, GSE36809 for human). To test the specificity of our cross-species transcriptional comparison, we utilized the mouse model for Duchenne’s muscular dystrophy (DMD, mdx mouse, GSE1025) as a biological negative control, and selected the peak pathological timepoints (Day 28) to benchmark our models. Detailed procedures regarding data normalization, probe selection, and quality control checks are provided in the **Supplementary Information**. Briefly, we evaluated the log_2_-expression and Log2FC distributions, filtered genes with low expression, performed PCAs to identify outliers and batch effects, and performed a differential gene expression analysis.

### 2.2. Conservation depends on statistical assumptions, stimulus intensity, kinetics and standard error

To evaluate the predictive validity of the mouse model across different immunological challenges, we analyzed the distribution of differentially expressed genes (DEGs) and the mean log_2_-fold values for all six conditions: Influenza (Fluad®) and Hepatitis B (Engerix **B**®) immunization, S. aureus and E. coli infections, and burn and trauma-hemorrhage (Trauma) injury. To enforce methodological symmetry, we implemented an iterative downsampling framework (subsampling human subjects to n = 5 over 200 iterations and summarizing statistics by the median), which effectively eliminated pseudo-significant inflation and established a reliable baseline for cross-species comparison (**Fig. 2a**).

**Fig. 2.**
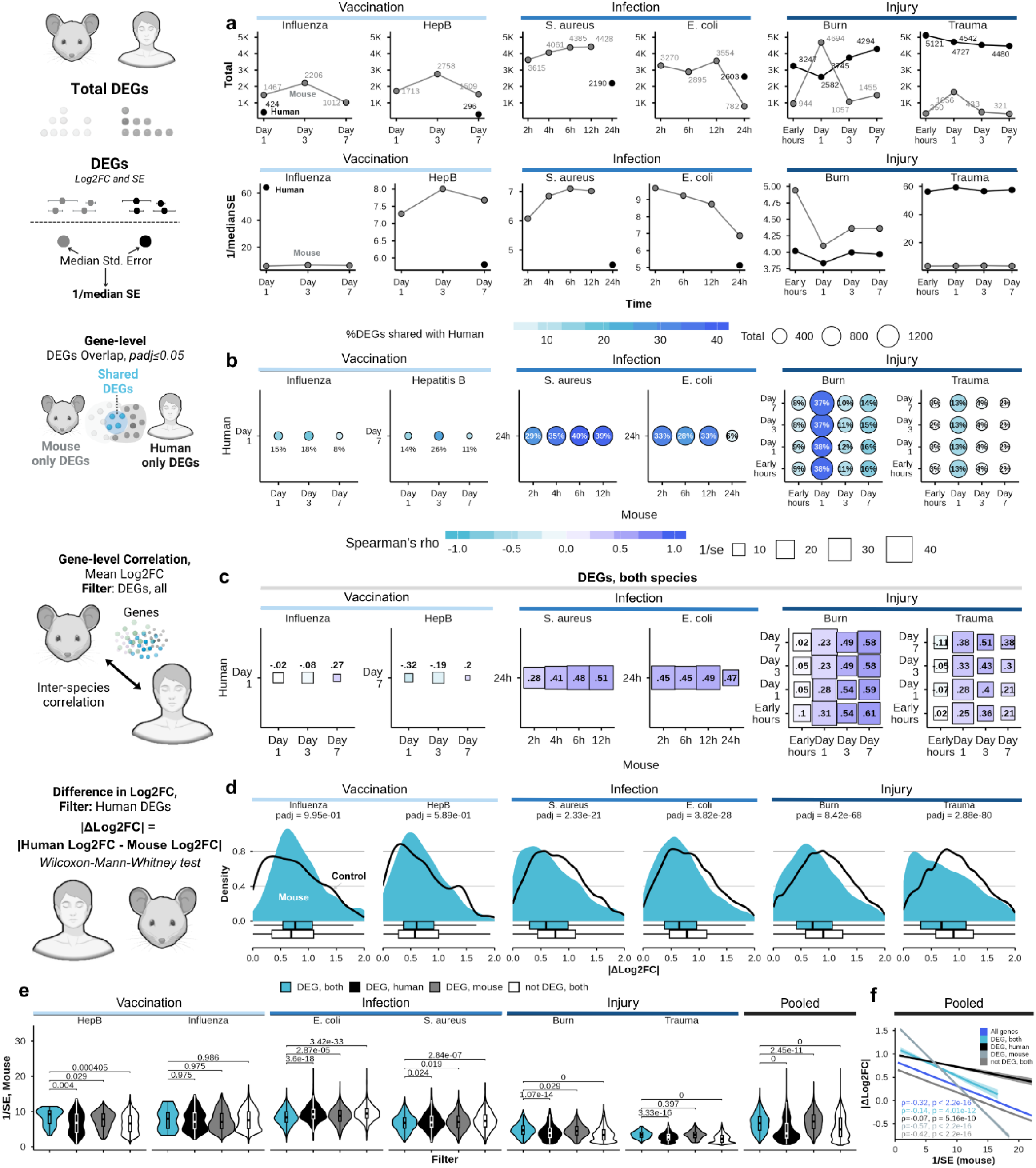
Gene-level conservation between murine and human expression. (a) Total number of DEGs (adjusted p < 0.05, top) and median inverse standard error (1/SE, bottom) across conditions. (b) Overlapping DEGs across time-point combinations, with bubble size proportional to the number of shared DEGs and colour to the percentage. (c) Weighted Spearman matrices of Log2FC between species, weighted by 1/SE. (d) Density of |ΔLog2FC| for human DEGs versus negative controls (WMW test). (e) Mouse 1/SE across conditions and pooled, by gene group, with WMW adjusted P values. (f) Linear regression of |ΔLog2FC| on 1/SE, stratified by gene overlap category.

To define the optimal temporal alignment between species, we compared the mean standard error of the effect estimates, the total and overlapping DEG counts, and the interspecies profile correlations across matched timepoints (**Fig. 2a–c**). Best-matched pairs combined low measurement error, high DEG overlap, and maximal interspecies correlation. Weighted Spearman correlations, using the inverse standard error (1/SE) as the weight, revealed stimulus-specific kinetic alignment, where murine timepoints sampled 2–6 h after challenge matched the human peak inflammatory window (days 1–3). This kinetic decoupling indicates that cross-species comparisons should be anchored to biologically equivalent timeframes rather than to chronological hours.

Translational predictability was governed primarily by stimulus intensity. Intense systemic perturbations, such as acute bacterial infection and severe injury, engaged highly conserved, evolutionarily stable transcriptional programs and achieved strong interspecies correlations (Spearman ρ = 0.41–0.59 for S. aureus, E. coli, and burn models; **Fig. 2c**). Consistently, human DEGs in these models showed significantly smaller absolute log2-fold-change differences (|ΔLog2FC|) relative to their mouse orthologs than did non-matching negative controls (Wilcoxon–Mann–Whitney [WMW] test, adjusted P < 1 × 10⁻¹⁵; **Fig. 2d**). By contrast, single-dose vaccination did not reach the systemic activation threshold required to engage core mammalian inflammatory cascades, yielding low correlations (Spearman ρ = −0.32 to 0.27) that remained near baseline controls.

Finally, linear regression showed that mouse 1/SE scaled inversely with cross-species expression divergence (|ΔLog2FC|) across all conditions evaluated, with condition-wise correlations ranging from Spearman ρ = −0.07 to −0.57 (P < 0.001; **Fig. 2e**). Mouse-only DEGs showed extreme divergence at low 1/SE, reflecting technical volatility, whereas shared DEGs (present in both species) retained higher 1/SE and converged toward minimal log2-fold-change divergence at high 1/SE (1/SE > 15). Together, these results indicate that 1/SE-weighted filtering suppresses technical noise and exposes a core of shared mammalian response genes.

### 2.3. Mouse expression classifies the direction of human DEGs

To quantify the predictive accuracy of murine transcriptional signatures in classifying human gene expression direction (up- vs. not-up regulated and down- not-down-regulated), we performed ROC curve analysis across conditions (**Fig. 3a**). Infection and injury models classified human gene regulation with ROC-AUC values ranging from 0.64 to 0.74 across full mouse gene sets, exceeding the biological negative control (DMD, ROC-AUC 0.48–0.61). Filtering for immune-related gene subsets yielded minimal gain overall, except for S. aureus infection (ROC-AUC values of 0.75 to 0.80). By contrast, vaccination models resulted in ROC-AUC values of 0.41 to 0.64, performing similarly to the DMD negative control and nearly random. Because class imbalance between DEGs and non-DEGs can skew ROC curves, we computed PR-AUC matrices to evaluate predictive robustness (**Fig. 3b**). The PR-AUC analyses corroborated the ROC findings, with infection and injury models achieving PR-AUC values of 0.53 to 0.76. In contrast, vaccination models exhibited PR-AUC similar to the negative control and with a small number of genes (n = 1 to 3). Across all evaluated conditions, predictive performance was asymmetrical between regulatory directions, with ROC-AUC and PR-AUC values for downregulated transcripts exceeding those for upregulated genes in early time points following infection and injury challenges.

**Fig. 3.**
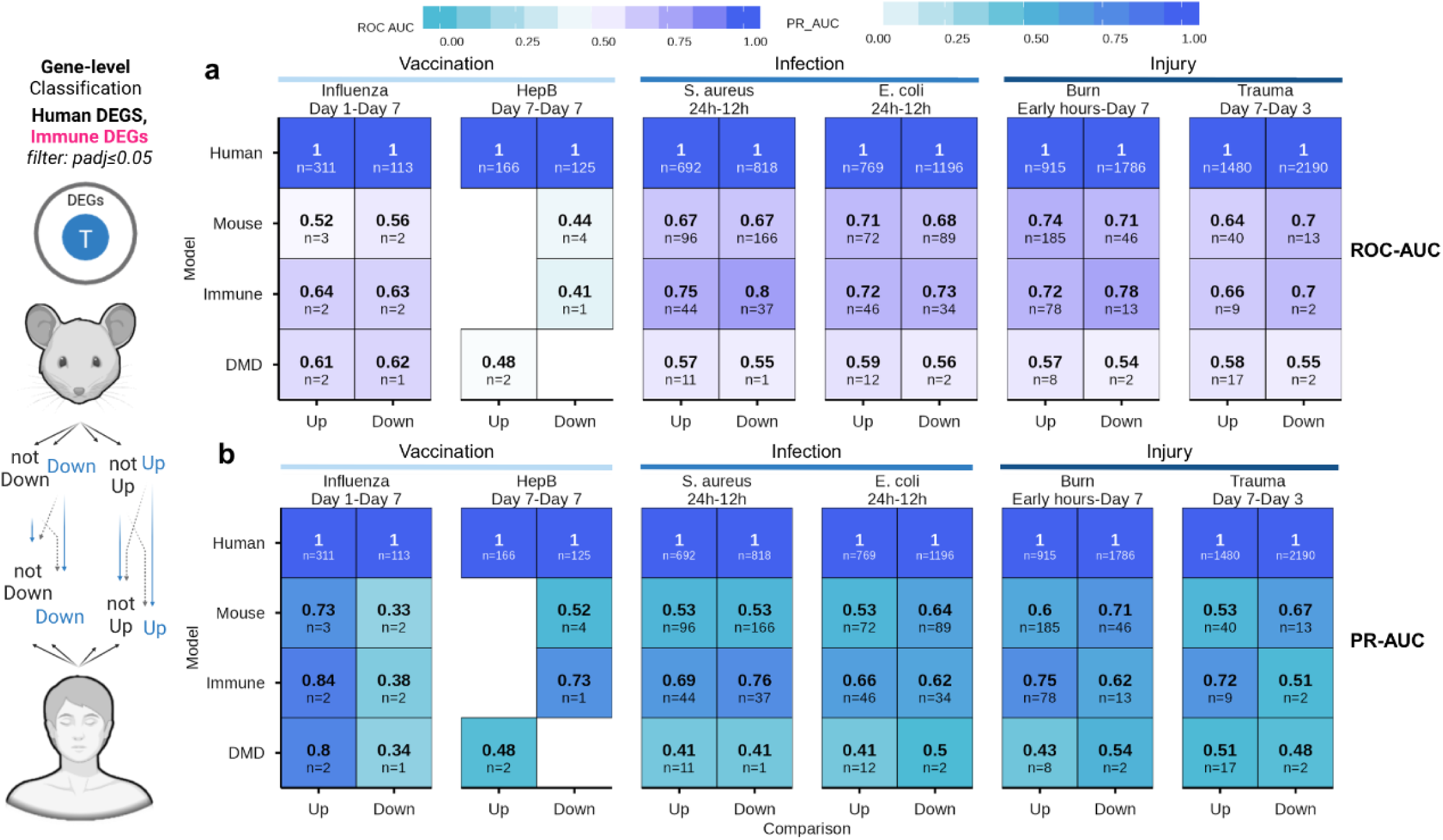
Gene-level classification performance across stimulus categories. (a) ROC-AUC matrices for mouse signatures classifying up- and downregulated human genes across vaccination, infection and injury. (b) PR-AUC matrices accounting for class imbalance. Both panels show full mouse gene sets (Mouse) and immune-related subsets (Immune), with human self-prediction as positive control and the DMD dataset as negative control.

We then used the prediction model FIT (Found In Translation), a lasso regression-based model trained on several cross-paired conditions including infections by influenza, *E. coli*, and *S. aureus,* and injuries by burn and trauma. When comparing gene expression in influenza vaccination between mouse and human, the FIT output was biased toward positive Log2FC (**Supplementary Fig. S3a**). We investigated the potential basis for this overprediction by analyzing the distribution of Log2FC for all curated gene sets in the training data. We found that the majority of conditions did not skew distributions, indicating this did not drive the prediction bias (**Supplementary Fig. S3b**). This discrepancy could also be explained by the differential expression analysis method used (z-test), which identified fewer DEGs than our analysis (**Supplementary Fig. S3c**), and the inclusion of different conditions and sample sources in the training dataset. Thus, we did not use this prediction tool further.

### 2.4. Modular analysis improves translational correlation and shows high sharing of leading-edge genes between species

To evaluate interspecies translatability at the pathway level, we analyzed immune responses using Blood Transcription Modules (BTMs). Module-level analysis using normalized enrichment scores (NES) showed higher interspecies correlation than individual gene-level analyses when we restricted the analysis to modules with adjusted P < 0.25 in mice or in both species (**Fig. 4a; Supplementary Fig. S4b**). Module-level correlations were higher in all conditions (Pearson r = 0.89 to 0.97) except trauma (Pearson r = −0.22 to 0.65, R² = 0.05 to 0.42; Fig. 4a; **Supplementary Fig. S4a**). As vaccinations had few significantly enriched modules (n < 5, adjusted P <0.25), we did not use these conditions further in our analyses. Furthermore, across all matched infection and injury conditions, except for trauma, mouse BTM models consistently outperformed DMD, which exhibited near-baseline or weak correlations (r = −0.32 to 0.28, P > 0.05; **Supplementary Fig. S6d**).

**Fig. 4.**
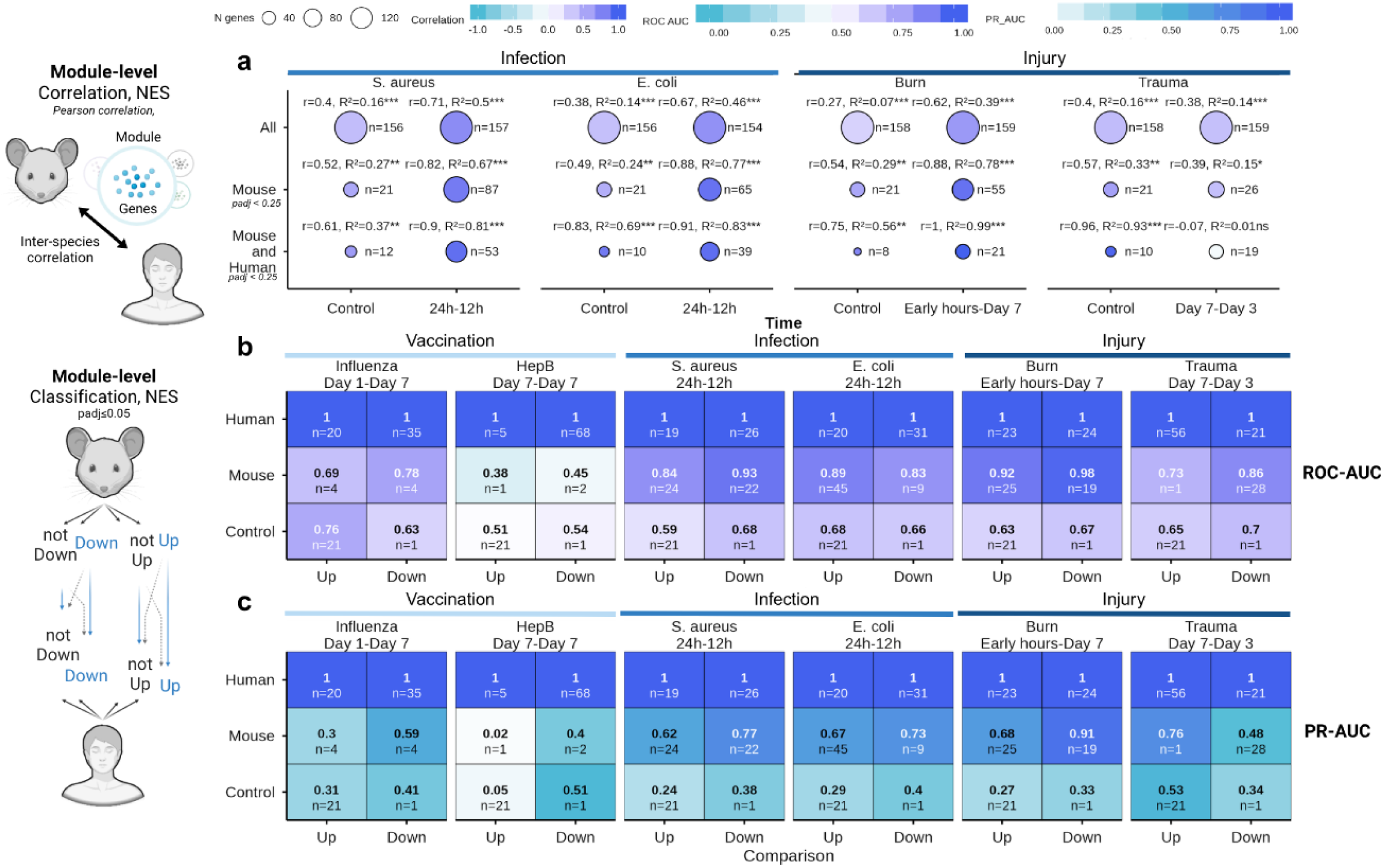
Immune response analysis using BTMs. (a) Pearson correlation of BTM NES between species at matched time points, for all enriched modules, modules enriched in mouse and modules enriched in both. (b) ROC-AUC of mouse-derived BTM enrichment signatures classifying human module regulation. (c) PR-AUC for the same predictions, comparing human, mouse and control datasets.

Classification performance based on mouse BTM enrichment signatures exceeded that of gene-level models in predicting human module regulation across infection and injury conditions. ROC-AUC values for module regulation ranged from 0.84 to 0.98 in infection and injury cohorts **(Fig. 4b**), and were markedly higher than those of DMD, which yielded near-random classification metrics (ROC-AUC 0.48 to 0.61; **Supplementary Fig. S6b**). PR-AUC analysis supported these findings, with mouse BTM models achieving values between 0.62 and 0.91 in acute challenges, compared with baseline control values of 0.24 to 0.53 (**Fig. 4c**).

To evaluate gene-level divergence within functional pathways, we analysed LEGs across BTMs (**Fig. 5**). The percentage of shared LEGs relative to the total number of mouse LEGs varied across stimulus conditions and process categories, with median values ranging from 67.5% to 79% in infection and injury conditions, and 70.5% when pooling all conditions **(Fig. 5a**).

**Fig. 5.**
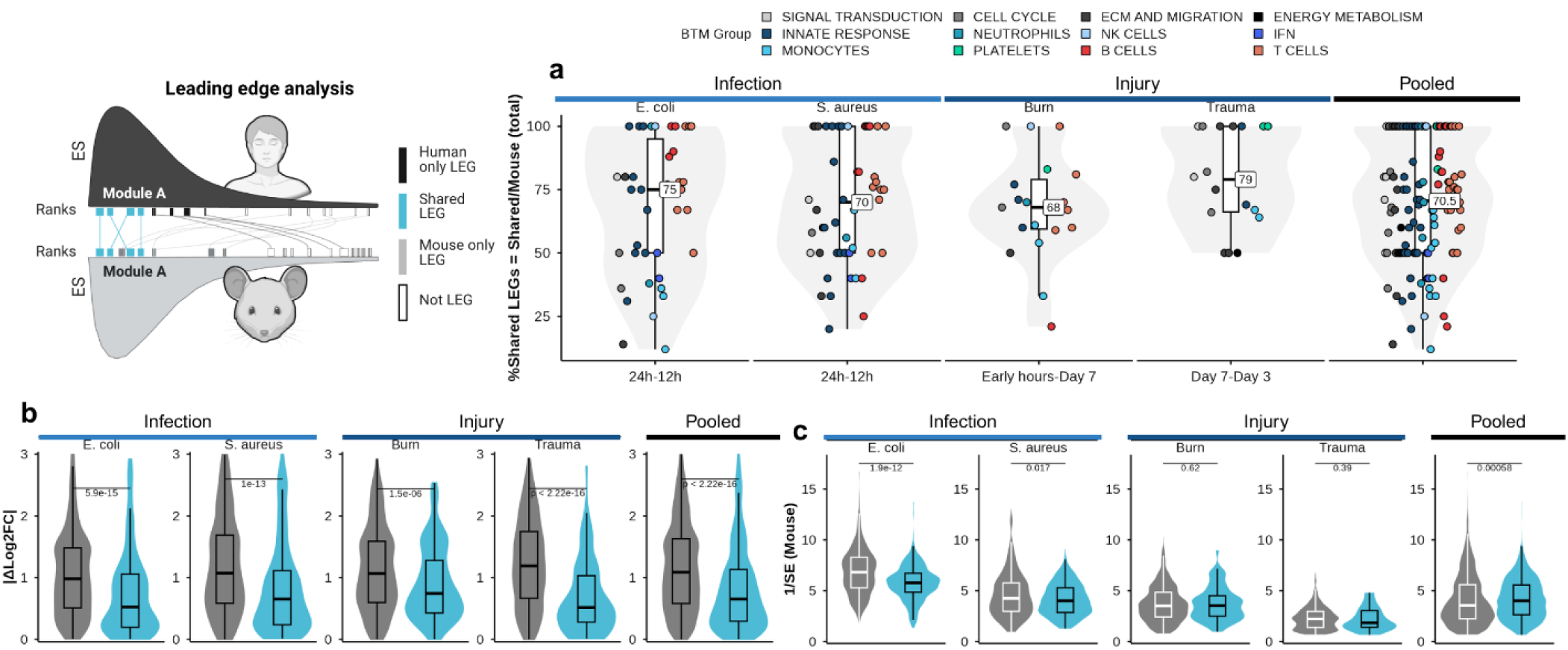
Leading-edge gene (LEG) analysis across BTMs. (a) Percentage of shared LEGs relative to total mouse LEGs across infection and injury, coloured and grouped by BTM process category. (b) |dLog2FC| between species across infection, injury and pooled datasets, with Wilcoxon P values comparing shared and non-shared LEGs. (c) Mouse 1/SE across the same conditions, compared between shared and non-shared LEGs.

We then asked why some LEGs are shared, and compared Shared and Not-Shared LEGs in terms of the absolute difference in log2 fold change (|ΔLog2FC|) and the mouse 1/SE across infection, injury, and pooled conditions. Overall, Shared LEGs showed a smaller |ΔLog2FC| (WMW, adjusted P < 2.22 × 10⁻16; **Fig. 5b**), but a lower 1/SE than Not-Shared LEGs (WMW, adjusted P < 0.00058; **Fig. 5c**). However, 1/SE did not differ significantly in burn and trauma (WMW, adjusted P = 0.62 and 0.39; **Fig. 5c**).

### 2.5. Gene sequence conservation and regulatory architecture are associated with cross-species expression divergence

To identify the molecular features associated with cross-species expression convergence, we evaluated sequence evolution and regulatory features across all high-confidence orthologs (**Fig. 6**). Continuous relationships between genomic/regulatory characteristics and cross-species expression divergence (|ΔLog2FC|) were evaluated using monotonic linear models and Spearman rank correlation coefficients (full regression equations, fitted slopes, and statistical parameters are reported in **Fig. 6**).

**Fig. 6.**
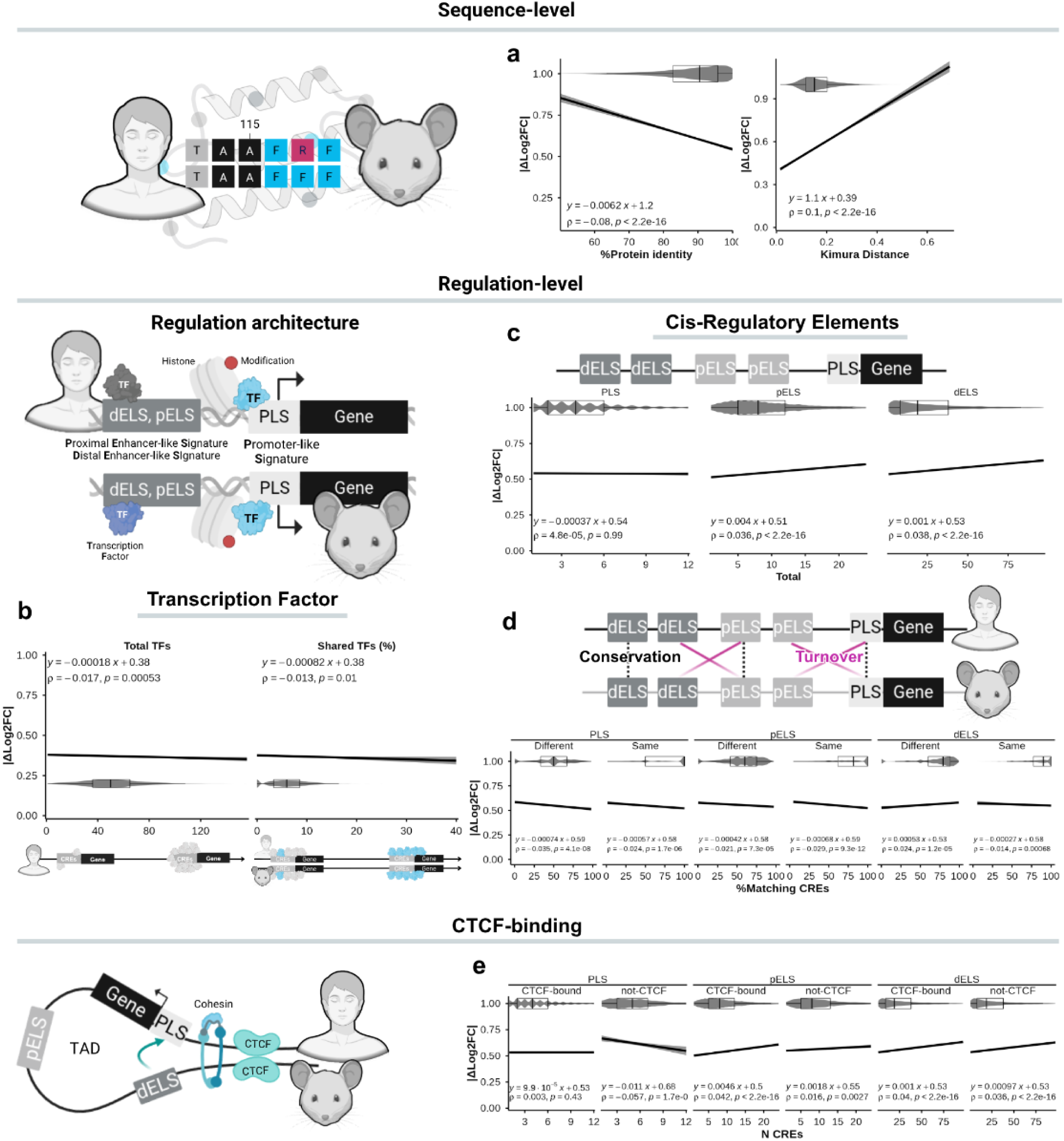
Impact of protein sequence identity, evolutionary distance, transcription factor binding and cis-regulatory element conservation on cross-species regulation. (a) Linear regression of |dLog2FC| on protein sequence identity or Kimura distance, stratified by shared LEGs, non-shared LEGs and all genes. (b) Expression alignment as a function of total human TFs (left) and the percentage of shared TFs (right). (c) |dLog2FC| versus total CRE counts for dELS, pELS and PLS. (d) Expression differences across CRE conservation and turnover states, with matching percentages compared across CRE types. (e) Distribution of |dLog2FC| across varying CRE numbers, by gene overlap category.

Protein sequence conservation was associated with reduced expression divergence. Across all high-confidence orthologs, higher sequence identity correlated with lower |ΔLog2FC| (Spearman rho = -0.08, P < 2.2 × 10-16; **Fig. 6a**), whereas greater Kimura evolutionary distances correlated with increased divergence (Spearman rho = 0.10, P < 2.2 × 10-16; **Fig. 6a**). Genes under stronger coding-sequence constraint therefore also tend to preserve their perturbation response magnitude between species.

At the trans-regulatory level, higher transcription factor (TF) availability was inversely associated with expression divergence. Both total human TF count (rho = −0.017, P = 0.00053; **Fig. 6b**) and the percentage of TFs shared between mouse and human (rho = −0.013, P = 0.010; **Fig. 6b**) negatively correlated with |ΔLog2FC|, indicating that orthologs with richer trans-regulatory architecture operate under tighter variance constraints.

Cis-regulatory element (CRE) multiplicity also modulated divergence across element classes (**Fig. 6c**). CRE abundance correlated positively with expression divergence for proximal enhancers (pELS: rho = 0.036, P < 2.2 × 10⁻¹⁶; **Fig. 6c)** and distal enhancers (dELS: rho = 0.039, P < 2.2 × 10⁻¹⁶; **Fig. 6c**), whereas promoter-like signatures showed no significant relationship (PLS: rho = 4.8 × 10⁻⁵, P = 0.99; **Fig. 6c**).

We next evaluated how CRE turnover impacts expression divergence by analyzing the percentage of matching elements between species (**Fig. 6d**). For pELS, turned-over elements (PLS - Different) showed a weak negative correlation with divergence (rho = -0.021, P = 0.00018; **Fig. 6d**), whereas conserved promoters (PLS - Same) showed no correlation (P = 0.38; **Fig. 6d)**. For pELS, matching CRE percentage negatively correlated with divergence regardless of turnover status (pELS - Different: rho = -0.038, P = 1.1 × 10⁻¹²; pELS - Same: rho = -0.028, P = 4.1 × 10⁻⁷; **Fig. 6d**). Conversely, dELS displayed positive associations across both categories (dELS - Different: rho = 0.012, P = 0.032; dELS - Same: rho = 0.038, P = 3.6 × 10⁻¹²; **Fig. 6d**).

Finally, CTCF binding altered the relationship between CRE multiplicity and divergence across classes (**Fig. 6e**). In pELS, CRE count increased divergence more strongly in CTCF-bound compared to non-CTCF-bound elements (bound: rho = 0.042, P = 2.3 × 10⁻¹⁰; non-bound: rho = 0.018, P = 0.0027; **Fig. 6e**). In contrast, dELS showed comparable positive associations regardless of CTCF binding (bound: rho = 0.04; non-bound: rho = 0.038; both P < 2.2 × 10⁻¹⁶; **Fig. 6e**), while PLS maintained weak or non-significant trends (**Fig. 6e**).

These associations describe divergence of magnitude. To test convergence while controlling for expression level, we next modelled rank-based targets.

### 2.6. Multilayer modelling shows that evolutionary and regulatory features explain cross-species differences

To quantify how evolutionary, regulatory and pathway features contribute to cross-species translatability, we constructed bottom-up models with sequential addition of two nested feature layers: a “DGE Baseline” containing only the mouse differential-expression summary (mouse rank for rank transfer; mouse direction for directional concordance; mouse 1/SE for shared-LEG classification), and a “Full + BTM” set that added the biological layers gene and protein sequence evolution, CRE architecture, and BTM group membership (**Fig. 7a**).

**Fig. 7.**
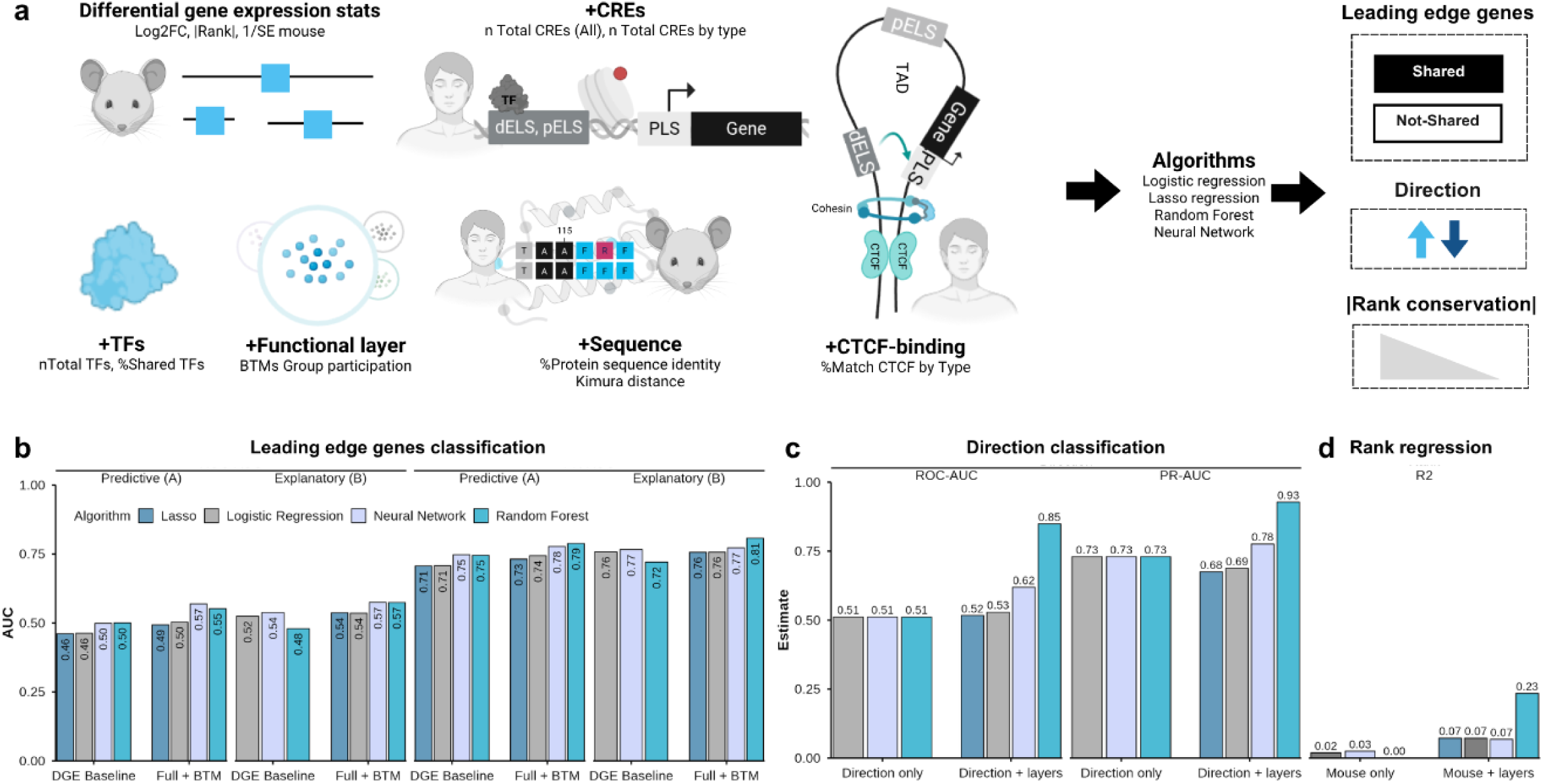
Multilayered predictive modelling connecting evolutionary, regulatory, and pathway features to translatability. (a) Bottom-up workflow of progressive layer integration, from the DGE baseline to sequence evolution, TF availability, CRE architecture, CTCF binding and BTM membership, evaluated by LOCO-CV (4 folds) across logistic regression, lasso, random forest and neural network. (b) Out-of-fold ROC-AUC for shared versus mouse-only LEG classification across model layers. (c) Out-of-fold ROC-AUC for directional concordance. (d) Out-of-fold R2 for prediction of the human absolute rank, |Log2FC| x -log10(adjusted P).

Two universes were used: all human DEGs (adjusted P ≤ 0.05) for rank transfer and direction, and mouse LEGs (Shared vs. Mouse only) for shared-LEG classification. Each gene received an absolute rank (0–100) from the percentile of the product of the absolute Log2FC and the −log10(adjusted P) within its universe and condition. Rank divergence was defined as the absolute difference between the mouse and human absolute ranks (lower values indicate greater convergence), and relevance as the lower absolute rank of the two species. Both metrics are rank-based and therefore direction-agnostic; directional concordance was evaluated separately as a distinct target. Models were evaluated by leave-one-condition-out cross-validation (four folds), comparing linear and lasso regression, random forest and a neural network (with logistic regression as the linear baseline for classification). To identify the primary molecular and statistical drivers behind these predictions, we extracted standardized feature importance metrics across both parametric (logistic/lasso regression) and non-parametric (random forest, neural network) architectures across all three modelling tasks (**Supplementary Fig. S7**).

Mouse ranks and directions did not transfer on their own: mouse rank alone explained almost no variance in human rank (R² = 0.001; linear 0.019; neural network 0.025), and mouse direction alone was near chance for directional concordance (ROC-AUC 0.511). Adding the biological layers improved both, raising random forest R² to 0.234 for human rank (lasso 0.072; linear 0.072; neural network 0.068) and random forest ROC-AUC to 0.849 for direction (neural network 0.619; logistic 0.529; lasso 0.517; **Fig. 7c,d**). For rank transfer, the most important random forest features were mouse rank, Kimura distance, total TFs and total CRE counts, which acted as positive drivers, together with protein identity, TF sharing and non-BTM membership, which acted as negative drivers (**Supplementary Fig. S7a,b**). For directional concordance, the most important random forest features were TF sharing, pELS CRE counts and pELS matching, which favoured concordance, together with Kimura distance, protein identity, total TFs and dELS CRE counts, which favoured non-concordance (**Supplementary Fig. S7c,d**). Sequence constraints and trans-regulatory availability therefore shape directionally synchronized responses during acute infection and injury.

Distinguishing shared from mouse-only LEGs was harder. Among mouse LEGs, the Lasso reached the highest out-of-fold ROC-AUC in both framings (0.617 in the predictive framing and 0.627 in the explanatory framing), marginally above the neural network (0.615 and 0.603), logistic regression (0.601 and 0.611) and random forest (0.572 and 0.622; **Fig. 7b**), and also retained the highest PR-AUC (0.915 and 0.917). Across the three tasks, the random forest was the best-performing algorithm for rank transfer and directional concordance, whereas the Lasso was marginally better for shared-LEG classification. The lasso improved only marginally over the unregularized logistic model for shared-LEG classification (0.617 vs 0.601) and not at all for rank transfer or direction. For shared-LEG classification, the lasso selected a sparse set of predictors, in which T-cell, innate-response, monocyte and B-cell module membership and the BTM-only mouse rank increased the odds of shared status, whereas higher dELS type matching, total TFs and PLS CTCF matching reduced it (**Supplementary Fig. S7e**). Results were unchanged when the analysis was repeated at the gene × module level, treating each module membership as a separate observation and including module identity as a predictor (ROC-AUC 0.621 in the predictive framing and 0.633 in the explanatory framing).

## 3. DISCUSSION

In this study, we show that murine predictability for human immunity is conditional on granularity and temporal alignment. By reconciling Seok et al. ^17^ and Takao & Miyakawa ^18^, we found that individual genes showed noisy linear correlation (Pearson r), whereas BTM activation and rank hierarchy (Spearman rho) were conserved. Unlike Seok et al., who mixed tissues in trauma/endotoxemia, we restricted the analysis to blood to limit tissue confounding.

We used limma with empirical Bayes shrinkage and covariate adjustment (age, sex, ethnicity), whereas Seok et al. used EDGE and Normand et al. ^22^ used a z-test. To address sample-size imbalance, human cohorts were downsampled and statistics summarized by the median. Compared with the z- and t-tests used previously, limma improves power, which contests prior reports of poor correlation. Therefore, much of the residual disagreement between studies is explained by the statistical test used. When the same adjusted-P threshold was applied, our gene-level correlations closely matched those of Takao and Miyakawa, although the identity and distribution of the selected genes differed. By contrast, Seok et al. retained far more DEGs than we did, even under a stringent threshold (adjusted P < 0.0001), likely because their EDGE spline-based pipeline was more permissive, and yet they reported gene-level Pearson correlations of only 0 to 0.09, well below the values we obtained when all genes were considered (Pearson r > 0.3, data not shown). Thus, the magnitude of the reported discrepancy depends jointly on the stringency of DEG selection and on whether the analysis is restricted to selected DEGs or spans the whole transcriptome.

Transcriptional magnitudes scale non-linearly between species, so we calculated a weighted Spearman correlation between human and mouse orthologous profiles. This rank-based approach is robust to outliers, and weighting by precision (1/SE) down-weights noisy gene pairs. Furthermore, optimal alignment rarely occurred at chronologically matched time points, indicating stimulus-specific kinetics. Future comparisons should therefore prioritize biologically equivalent over chronologically matched time points.

Our evaluation of the FIT model revealed a systematic positive bias toward up-regulated genes, limiting accuracy for human responses. This benchmark is limited to a single model trained on a different pipeline (z-test vs. limma), heterogeneous tissues (liver, spleen, lung and brain vs. peripheral blood) and chronic conditions (e.g., rheumatoid arthritis, inflammatory bowel disease and Alzheimer’s disease vs. acute responses). A systematic comparison of algorithms trained exclusively on blood-derived cross-species data is beyond the present scope but represents a clear direction for future work.

Beyond statistical refinement, translatability was dictated by the nature and magnitude of the perturbation. Intense systemic perturbations, such as acute bacterial infections (S. aureus, E. coli) and severe tissue trauma (burn and haemorrhage), induced conserved transcriptional programmes, reaching high predictive accuracy at both gene and module levels (Fig. 3 and Fig. 4). By contrast, milder perturbations, such as single-dose subunit or inactivated vaccines (Fluad ® and Engerix B ®), showed species-specific divergence and lower predictive values. Single-dose immunizations likely fail to reach the activation threshold for core mammalian inflammatory cascades, instead exposing regulatory differences in early kinetics and baseline cell composition. Therefore, preclinical vaccine evaluation may therefore require booster regimens or potent adjuvants to surpass species-specific noise. Additionally, strain background matters, as C57BL/6 mice are Th1-biased and may limit predictability for Th2-dominated responses, requiring alignment of strain choice with the immunological hypothesis ^27^.

At the pathway level, collapsing high-dimensional transcriptomic data into functional modules such as BTMs ^28^ improved cross-species concordance. GSEA, which scores the full ranked distribution without arbitrary thresholds, yielded higher and significant correlations, with mouse-enriched modules correlating with human counterparts. The main translational signal lay in LEGs. Because non-responsive genes dilute pathway-level correlations, isolating LEGs captured the drivers of immune activation. Shared LEGs showed higher 1/SE in mouse, correlating with lower |ΔLog2FC|.

In addition to methodological issues, the lack of direct linear correlations in individual genes may be linked to species-specific differences in gene sequence evolution, regulation, and technical noise. However, evolutionary biology suggests that while individual gene sequences and regulation may diverge, the underlying biological processes tend to be conserved. Our results support this, because by collapsing data dimensionality into modules (BTMs), we reduced noise and revealed a robust translational signature.

To understand why specific orthologs maintain expression magnitude under perturbations while others diverge, particularly for genes with high absolute ranks (strongly up- or down-regulated), we integrated coding sequence evolution, regulatory architecture conservation, and functional attributes for each gene. Protein evolution and regulatory conservation jointly shaped translatability, indicating that translatability extends beyond purely regulatory or purely transcriptomic models. While higher protein sequence identity and lower Kimura evolutionary distances correlate with reduced expression divergence (|ΔLog2FC|) between species, non-coding regulatory elements undergo substantial evolutionary turnover ^7^. Although individual transcription factor binding site occupancies diverge rapidly across mammalian lineages, higher-level trans-acting regulatory networks and chromatin domain structures remain broadly conserved ^29^. Accordingly, genes that undergo CRE turnover or display divergent CTCF chromatin insulation exhibit heightened expression divergence (|ΔLog2FC|) between humans and mice. Consequently, cross-species divergence in immune gene expression reflects a composite architecture governed simultaneously by coding sequence constraint and the dynamic structural remodelling of non-coding regulatory landscapes ^30,31,14,29^.

Our multilayer models showed that mouse rank and direction did not predict human responses on their own. Mouse and human ranks were only weakly correlated, mouse rank alone explained almost none of the variance in human rank, and mouse direction alone was only marginally better than chance at predicting directional concordance. Adding evolutionary and regulatory layers provided a conserved signal, with a marked increase in both random forest R² for human rank and ROC-AUC for directional concordance (**Fig. 7c,d**). The ranking of the algorithms, and the finding that the transferable signal is predominantly non-linear and multivariate, is reported in Section 2.6.

Relevance and convergence are distinct axes, and distinguishing shared from mouse-only LEGs is difficult. Among mouse leading-edge genes, classification performance was weak, showing that being a mouse LEG does not determine being a shared LEG, even though shared LEGs account for 67.5 to 79% of mouse LEGs in statistically significant enriched modules. The same biological layers were, however, decisive for rank transfer and directional prediction.

Beyond methodological advances, the real-world translation of preclinical models is frequently hindered by barriers to computational accessibility and reproducibility. Previous cross-species prediction tools were built on proprietary software environments such as MATLAB ³², while FIT suffers from an inactive web server and a GitHub repository that lacks data processing pipelines to independently reproduce or retrain the model²². Consequently, independently auditing these algorithms, applying them to new immunogenic perturbations, or retraining them with custom multi-omic layers remains challenging. In contrast, our framework prioritizes open science and practical usability. We provide step-by-step instructions for downloading and processing raw datasets, together with an R Markdown notebook that applies the trained models and computes the four translational scores for candidate prioritization. By releasing the complete open-source codebase, raw-data processing pipelines and trained models in a public repository, we empower the scientific community to audit, adapt and continuously retrain the architecture with additional perturbation models, booster vaccine regimens and emerging multi-omic layers.

### Limitations

Several limitations of the current study must be acknowledged. First, murine cohorts were small, which constrained statistical power, and the scarcity of datasets with sufficient metadata limited generalizability.

Regarding genetic diversity, the analysis was restricted to two inbred strains (C57BL/6 and CB6F1), which narrowed the phenotypic heterogeneity captured. Strain choice also matters, because host genetics shapes translatability. Analyses relied primarily on Th1-biased C57BL/6, in contrast to Th2-biased BALB/c. Low correlation in some modules therefore reflects strain–hypothesis mismatch rather than species-level failure, underscoring the need to align strain with the immunological hypothesis.

Beyond genetics, biological and experimental mismatches posed challenges. Laboratory mice are immunologically naïve, whereas humans carry complex exposure histories, a difference evident in Fluad® data. Mouse vaccinations were single-dose, whereas human inactivated/subunit vaccines typically use prime-boost, limiting comparison of adaptive trajectories. Human burn/trauma datasets also lacked standardized severity scores, precluding adjustment for this confounder.

At the genomic level, analyses relied on Ensembl-annotated one-to-one orthologs. The 50% amino-acid identity baseline established evolutionary relatedness but does not capture domain topology conservation ^33^, rewiring of protein–protein interactions across pathways ^34^, or the distinction between neutral and functionally impactful mutations.

Finally, regional constraint (PhastCons/phyloP), systematic transposable-element mapping, and expression of associated TFs were not included, and confounders such as baseline expression, gene length, GC content, CRE sequence alignment, and substitution-rate acceleration were not controlled. Scores were derived from blood-based BTMs and require testing in other tissues and gene sets (e.g., MSigDB, Gene Ontology).

## CONCLUSION

This study shows that murine translatability in systems immunology and vaccinology depends on analytical granularity and temporal alignment. Shifting from gene-centric to module-level analysis reduced technical noise and revealed conserved pathway activation, with activation hierarchy and mean module magnitude outperforming isolated orthologs as predictors. Accuracy was dictated by stimulus intensity, as acute infections and injuries engaged conserved profiles with higher concordance than single-dose inactivated/subunit vaccination.

Adding the biological layers (sequence evolution, TF features, CRE architecture, CTCF status and BTM membership) to the mouse DGE baseline improved prediction across all tasks. The random forest was the best-performing algorithm for human rank transfer and directional concordance, whereas a lasso-penalised logistic model was marginally better for shared-LEG classification by ROC-AUC (0.617 and 0.627 in the predictive and explanatory framings, respectively). The lasso improved only marginally over the unregularized linear model for shared-LEG classification and not at all for rank transfer or direction, indicating that the signal for rank and direction is predominantly non-linear and multivariate, while LEG sharing is largely captured by a linear, modular signal. The modular layers were essential, as mouse-only features reduced every algorithm to near chance for direction, whereas adding the BTM context raised the random forest to R² = 0.23 for human rank and ROC-AUC = 0.85 for directional concordance. The resulting out-of-fold predictions yielded four complementary scores, namely a sharing probability, a predicted human rank, a direction probability and a combined translational score. These scores are implemented in a step-by-step R Markdown notebook that applies the trained models to new murine datasets.

Future work should test prime-boost and adjuvanted regimens for low-immunogenicity platforms, refine ortholog regulatory annotation and standardize clinical metadata to delineate murine predictability. Designs should align strain with the hypothesis and match time points by biological equivalence rather than chronological time. Validation beyond BTMs is required across tissues and MSigDB/GO sets, with systematic TE mapping and PhastCons/phyloP scores.

## METHODS

### Data acquisition

To curate the mouse datasets, a structured filtering strategy was applied using the NCBI BioProject and the GEO databases. In BioProject, the initial search combined vaccine-related terms (“vaccine,” “vaccinated,” “vaccination,” and “vaccines”) with the filters “transcriptome gene expression,” “material transcriptome,” “capture_whole,” and “org mammals,” while excluding “*Homo sapiens*” to focus on non-human mammalian studies. A subsequent query was performed to isolate studies specifically related to *Mus musculus* by incorporating the organism filter and excluding records associated with tumors, cancer, and autoimmune diseases. Additionally, only datasets labeled under “Expression profiling by array” or “Expression profiling by high throughput sequencing” were retained.

Studies were manually annotated, and only those involving vaccines against human pathogens were included. To complete the annotation process, BioProject records were merged with GEO entries using PRJNA accession IDs. Each entry was then split to assign unique identifiers based on the vaccine used, sample source, and RNA-sequencing protocol. Datasets were excluded if they contained only partial methodological information or focused solely on BCR/TCR repertoire data.

Studies were annotated using GEOquery and then manually curated using information from the BioProject record and the corresponding published article. Furthermore, we annotated studies with details such as sample source, target condition and antigen, vaccine type and adjuvant, time points, doses, administration route, RNA assay method, and whether they were available on the 13Vax/covid-19Vax atlas or the MSigDB Vax collection. In cases where information conflicted with the paper, we prioritized the data present in the GEO records, such as the immunization route and the sample tissue source.

Clinical metadata were curated and standardized to ensure consistency across the dataset. For the human injury datasets, the time elapsed since injury, originally recorded in hours, was converted to days. To account for the irregular temporal distribution of samples, time points were discretized into categorical bins to align with the mouse cohort sample collection time points: <1 day (Early Hours), 1–2 days (Day 1), 2–5 days (Day 3), 5–10 days (Day 7), 10–18 days (Day 14), 18–25 days (Day 21), and >25 days. Samples were stratified into “Trauma” or “Burn” and “Control” groups based on subject identifiers.

### Preprocessing and quality control

In microarray datasets, probes mapping to the same Entrez ID were collapsed by selecting the one with the highest median expression. To map orthologous genes in humans, we used the protein-coding genes that have one-to-one orthology with human genes provided by the MGI database. Quality control was performed using the ArrayQualityMetrics package and limma diagnostic plots (**see Supplementary Information section)** ^35^.

### Differential gene expression and functional analysis

Expression data were log2-transformed, and low-abundance genes were filtered out, retaining only those with a log2-intensity higher than 6 in at least five samples to reduce technical noise and improve statistical power. For the E. coli mouse dataset, weight estimation was stratified by experimental batch (var.group = batch) to mitigate batch effects while preserving the biological signal of the infection. For longitudinal data, we treated individuals as random effects, and for both longitudinal and case-control human datasets, we included other covariates such as sex, ethnicity, and age as fixed effects in the limma matrix design.

To ensure robust detection of shared transcriptional signatures, we conducted differential expression analysis using the limma package and contrasted its performance with conventional metrics used in prior studies, such as standard t-tests ^36,37^ (Fig. S2). By employing empirical Bayes variance shrinkage and adjusting for demographic covariates and batch effects, the limma framework significantly outperformed the standard paired t-test in our datasets. Specifically, the t-test approach failed to identify significant DEGs in the vaccinated mouse cohorts, which would have led to the erroneous conclusion of limited cross-species conservation. To account for heteroscedasticity across clinical and experimental samples, sample-quality weights were estimated using the arrayWeights function, ensuring that high-variance samples had a reduced impact on the linear model. The trend and robust parameters were applied to account for the mean-variance relationship and minimize the influence of outliers.

To account for sample size imbalances between human and mouse studies, we downsampled human samples to 5, with bootstrap over 200 iterations, and used the median value for statistics derived from limma. Genes with an adjusted p-value smaller than 0.05 were defined as DEGs.

### Correlation and functional enrichment analysis

Correlation tests were performed at both the gene and modular levels. Gene-level cross-species comparisons were evaluated using weighted Spearman rank correlations (rho), with weights defined as the inverse standard error (1/SE) of the murine effect size estimates. Module-level correlations used Pearson correlation weighted by −log10(adjusted P) from GSEA. At the module-level, gene set enrichment analysis (GSEA) was first performed using the clusterProfiler ^38,39^, where genes were pre-ranked by the product of log2-fold change and -log10(p-adj), The BTM dataset was retrieved from a previous publication^28^. Module enrichment concordance was then evaluated using Pearson correlation coefficients (r) on Normalized Enrichment Scores (NES).

### Model evaluation

Model performance was evaluated using receiver operating characteristic (ROC) analysis to assess the ability to classify up- and down-regulated genes. Because limma produces two-sided p-values, one-sided p-values were recalculated from the moderated t-statistics by considering the upper tail for upregulated genes (log₂FC > 0) and the lower tail for downregulated genes (log₂FC < 0). The resulting one-sided p-values were subsequently adjusted for multiple hypothesis testing using the Benjamini–Hochberg false discovery rate correction.

As a reference, the corresponding human dataset was used as the reference “truth”, while predictions from the mouse model were compared against it. Two control models were also generated: a “perfect” model, which exactly replicated the human truth labels, and a “negative control” model, which used the DMD gene expression table. . For each gene and timepoint, prediction probabilities were defined as (1 - adjusted one-sided p-value), and ROC and PR curves were constructed. The area under the curve (AUC) was then computed to measure predictive performance. The same was applied to the functional modules, using the NES and the -log_10_(p-adjusted) derived from GSEA. This ROC-based framework evaluates module-level transfer (**Fig. 4b).**

### Gene sequence evolution and regulation architecture

Coding sequences (CDS) and amino-acid sequences for human–mouse protein-coding genes with confidence score = 1 were obtained from Ensembl via biomaRt, retaining Ensembl protein identity values. For Kimura distance ^40^, the longest CDS per gene was retained, aligned with pwalign ^41^ and scored as Kimura 2-parameter (K80) distance with ape ^42^.

To interrogate promoter and enhancer rewiring, cCREs were integrated from the ENCODE project ^43^ via the SCREEN portal, where the complete human (GRCh38) and mouse (mm10) cCRE catalogs and homologous cCREs were downloaded. We filtered for human cCREs categorized as PLS, pELS, or dELS, classified by whether they were CTCF-bound or non-bound. These cCREs were assigned to their associated human genes and subsequently compared to the annotated murine cCREs. Finally, we calculated the total number of cCREs that matched between species in terms of type (PLS, pELS, and dELS) and CTCF binding status, determining the corresponding percentage for each category.

Continuous relationships between sequence/regulatory features (e.g., sequence identity, Kimura distance, TF counts, CRE multiplicities) and expression divergence (|ΔLog2FC|) were fitted using ordinary least-squares (OLS) linear models and formally tested using monotonic Spearman rank correlation coefficients to account for non-normality and heteroscedasticity.

### Multilayer machine-learning modelling

To integrate evolutionary, regulatory and transcriptomic metrics, we assembled a gene-wise feature table comprising mouse log2FC, 1/SE, mouse absolute rank (overall and within BTMs), protein identity, CDS (Kimura) distance, total human TFs, mouse–human TF sharing, CRE architecture and BTM membership. Models were evaluated under leave-one-condition-out (LOCO) cross-validation across the four infection/injury folds (S. aureus, E. coli, burn, trauma), so that every gene is predicted by a model trained without its condition.

Two nested feature layers were compared, namely a DGE Baseline containing only the mouse differential-expression summary and a Full + BTM set that added the biological layers (sequence evolution, TF features, CRE architecture and BTM membership). Three tasks were modelled, namely shared-LEG classification (Shared vs Mouse only), human rank transfer (predicted human absolute rank, 0–100) and directional concordance (same vs opposite sign). For each gene and condition, rank convergence was defined as the absolute difference between the mouse and human absolute ranks, and relevance as the lower absolute rank of the two species. Because both metrics are rank-based and therefore direction-agnostic, directional concordance was modelled as a separate target. Two complementary scores were derived. The first is an observed prioritisation score, which multiplies cross-species convergence by the observed directional concordance and ranks genes measured in both species. The second is a predictive score (score_translational, defined below), which multiplies the predicted human rank by the predicted concordance probability and forecasts the human response for new murine data. The two scores therefore capture different quantities by design: observed cross-species agreement versus predicted human response magnitude.

Four algorithms were benchmarked under identical recipes, namely linear/logistic regression, lasso, random forest and a single-hidden-layer neural network, all implemented in tidymodels ^44^. Feature importance was assessed by log-odds (logistic), standardized effects (linear), lasso (penalized) coefficients and Gini importance (random forest). The random forest was the best model for human rank transfer and directional concordance, whereas the lasso was marginally better for shared-LEG classification by ROC-AUC; the lasso improved only marginally over the unregularized linear model for shared-LEG classification and not at all for rank transfer or direction.

Scores were derived from the out-of-fold predictions of the best model per task, so that every gene is scored by a model trained without its condition. These scores are score_shared (probability of being a shared LEG), score_rank (predicted human absolute rank, 0–100), score_direction (probability of concordant direction) and score_translational (score_rank × score_direction). Performance was measured by ROC-AUC and PR-AUC for classification and by R² and RMSE for regression.

To make the workflow reusable, we provide a step-by-step R Markdown notebook that applies the trained models to new murine datasets.

## Data availability

All source code, dataset-processing scripts and model-training pipelines are available at https://github.com/wapsyed/mousetohuman_multilayer. The step-by-step prediction notebook and the extensible model-building framework, which allow users to train customized cross-species models, are provided in the companion repository https://github.com/wapsyed/mousetohuman_predict.

## Author Contributions

WAPS – Conceptualization, coding, and manuscript writing; AAL, NC, EC, JDQS, BH, RDC – Manuscript contributions;

GC-M, TH, JEK, OCM, ECS – Supervision, editing, and approval of the final version.

## SUPPLEMENTARY INFORMATION

### Data preprocessing

For normalization, we selected the probe with the highest median expression for each gene among those sharing the same Entrez ID and then selected the Entrez IDs used by FIT. However, for normalization with human genes, we used the Entrez ID with the highest median expression as the reference for each gene when some genes had different Entrez IDs. We excluded genes that had multiple orthologs. These genes, identified by Entrez ID, were used as input for the FIT model, along with their corresponding log_2_-fold changes (L2FC), derived from a baseline contrast against pre-vaccination samples (0 hour).

To evaluate the quality of log_2_-transformed data, we generated density plots, boxplots, scatterplots, and performed PCA. A significant separation of a subset of human trauma samples was identified and excluded, as the corresponding metadata did not help to identify the cause by any covariate. However, we hypothesize that this could be explained by the severity of the trauma, which was not reported.

To evaluate the statistical integrity of our differential expression analysis, we inspected the raw p-value distributions across all comparisons. For most conditions, the histograms exhibited a characteristic enrichment of low p-values (p < 0.05). In specific cases where the adjusted p-value yielded fewer significant genes, such as in certain murine injury models, the raw p-value histograms revealed a lack of the expected peak at zero or a conservative upward trend, suggesting lower statistical power or high intra-group variability. The Benjamini-Hochberg correction was systematically applied to all contrasts to maintain a stringent False Discovery Rate (FDR) across the study.

Despite the implementation of advanced linear modeling techniques, the murine Burn and Trauma datasets exhibited near-uniform adjusted p-value distributions. This pattern reflects the high inter-individual biological variance inherent to cross-sectional studies compared to the longitudinal designs used in our other cohorts.

## SUPPLEMENTARY TABLES

**Supplementary Fig. S1.**
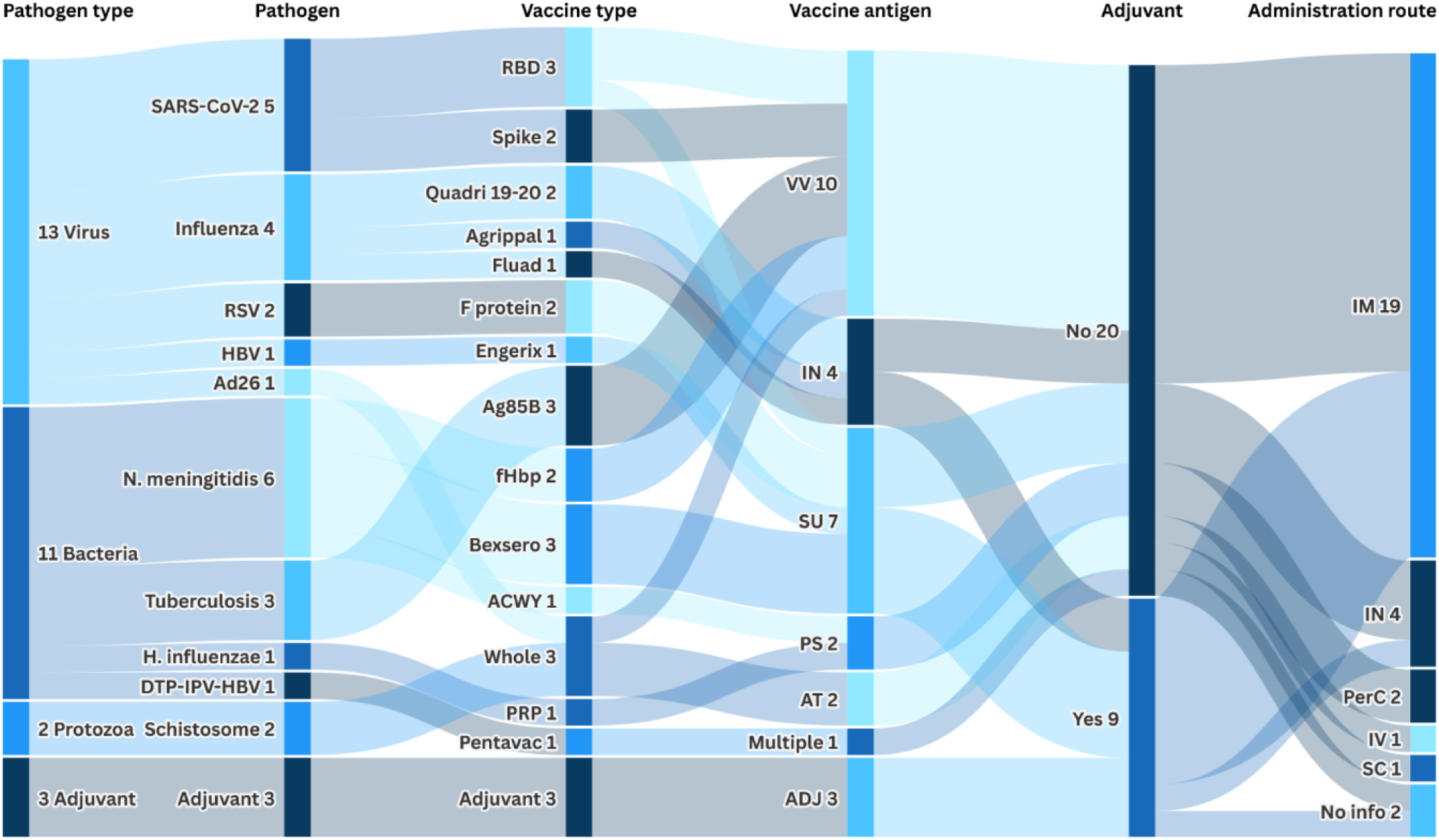
Sankey diagram of dataset distribution and study characteristics, across six categorical dimensions. Link width is proportional to the number of datasets. Categories are pathogen type (viral, bacterial, protozoan), pathogen or vaccine (for example Ag85B, spike protein, F protein), vaccine type (SU, subunit; VV, viral vector; IN, inactivated; PS, polysaccharide), adjuvant (yes or no) and administration route (IM, IN, PerC, IV, SC).

**Supplementary Fig. S2.**
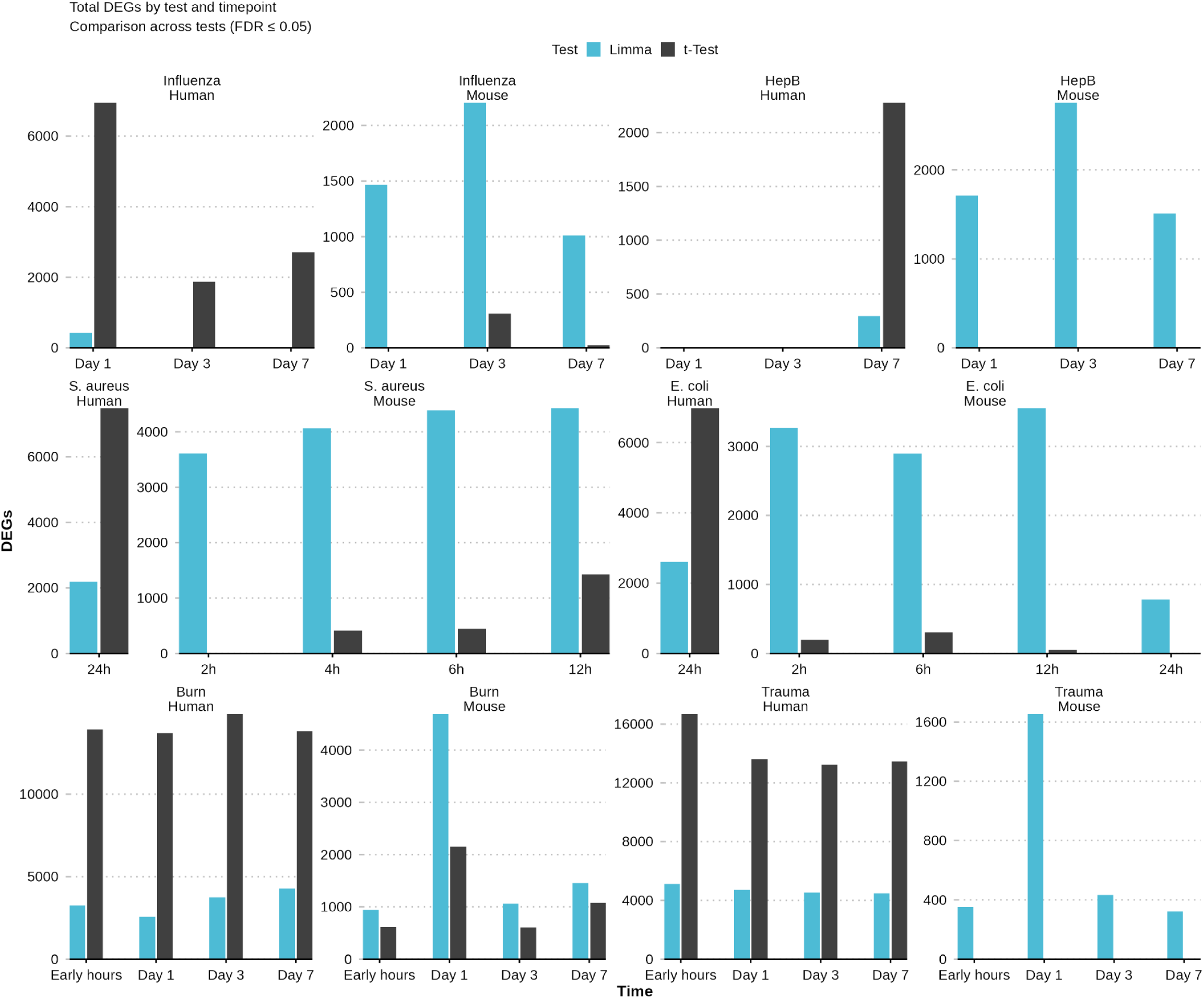
Comparison of statistical methodologies for Differential Gene Expression (DGE) analysis across species. Bar charts show the number of DEGs identified in human and mouse datasets across four conditions (Fluad, hepatitis B, S. aureus, E. coli), comparing limma (blue) and standard t-test (grey) and paired (dark) versus unpaired (light) designs. The y-axis is the total DEG count per time point. The comparison highlights the sensitivity of limma in paired clinical designs.

**Supplementary Fig. S3.**
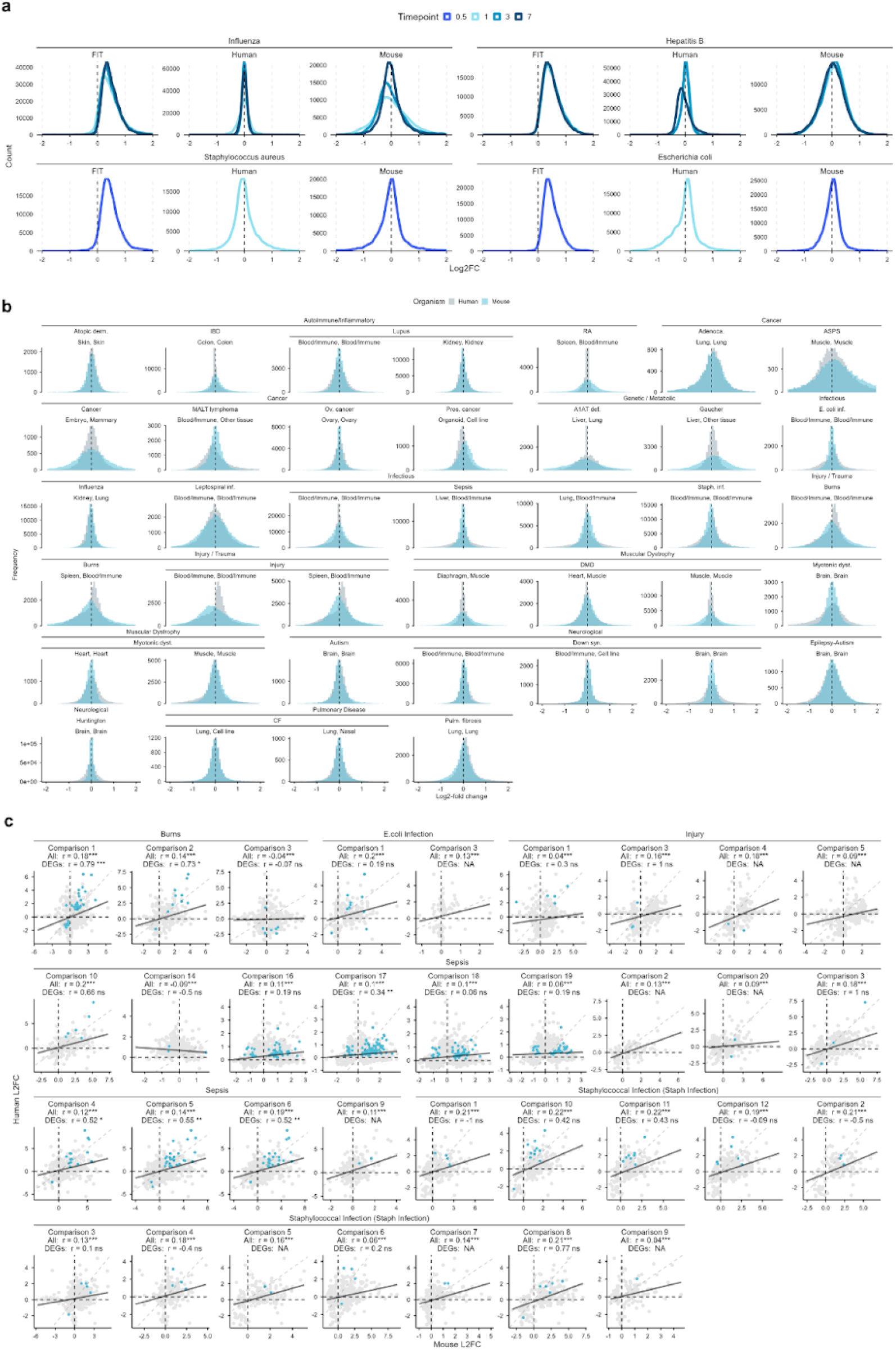
Cross-species distribution and correlation of gene expression changes in the FIT training dataset. Overlaid histograms show log2-fold changes (L2FC) in human and mouse samples across the diseases and tissues included (left). Scatter plots show the relationship between mouse and human log2FC for blood-related comparisons. Not-DEGs are in grey, and DEGs in both species (FDR-adjusted q <= 0.05) in blue. Spearman coefficients are reported for all genes and for shared DEGs. Asterisks indicate significance (*p < 0.10; **p < 0.05; ***p < 0.01). Correlations used comparisons with at least three shared observations. Disease abbreviations and group assignments are given in the associated metadata tables.

**Supplementary Fig. S4.**
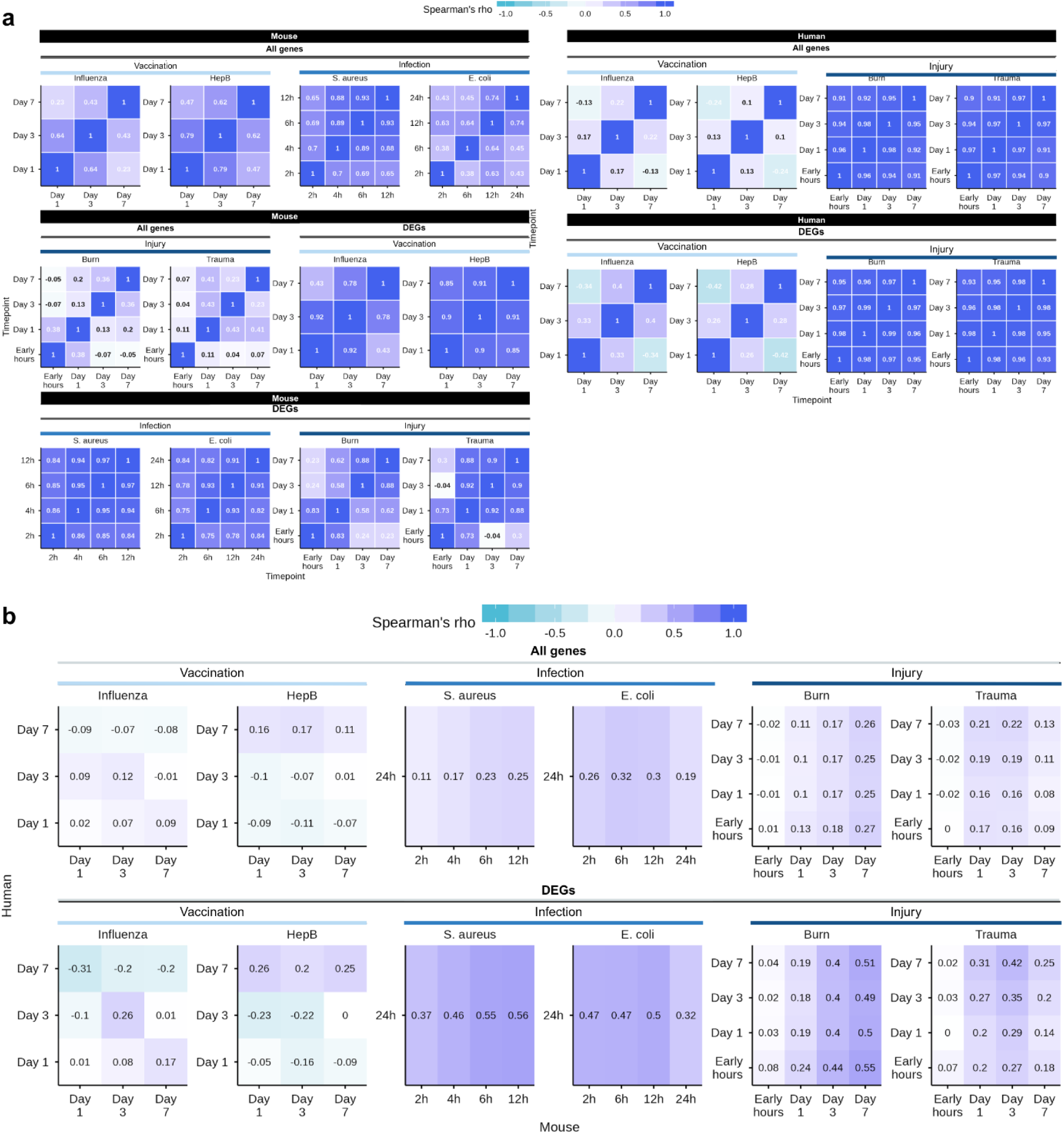
Correlation matrices of intra-species (a) and inter-species comparisons (b). Spearman rho was calculated from the Log2F of each species over time.

**Supplementary Fig. S5.**
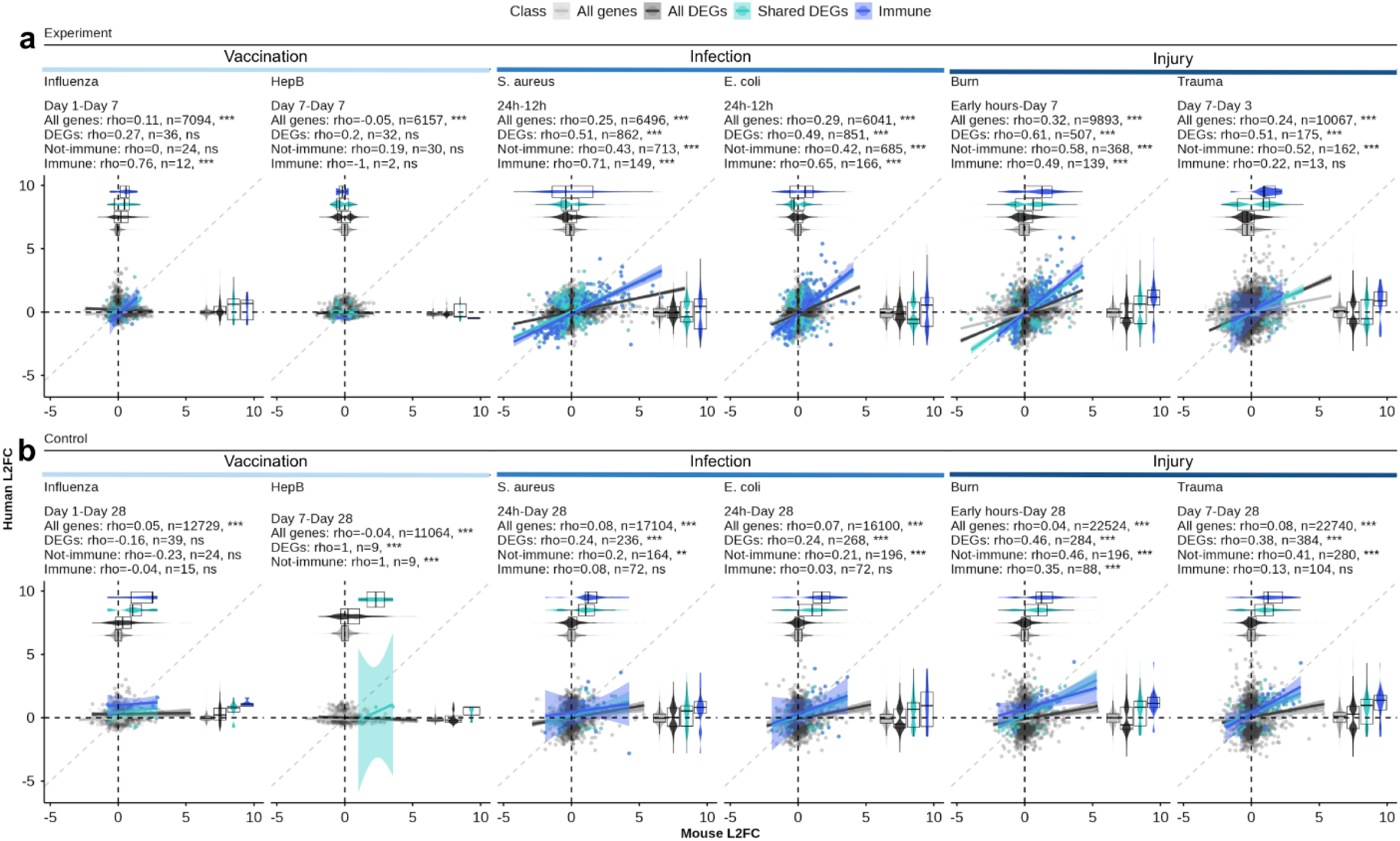
Gene-level correlation between humans and mice. Scatter plots illustrating the correlation of log_2_-fold change (log2FC) between human (y-axis) and mouse (x-axis) orthologous genes. Each dot represents a DEG in either species; colored dots represent DEGs in both species (blue), and immune DEGs (cyan). Spearman’s rank correlation coefficient (r) and sample size (n) are indicated for each comparison.

**Supplementary Fig. S6.**
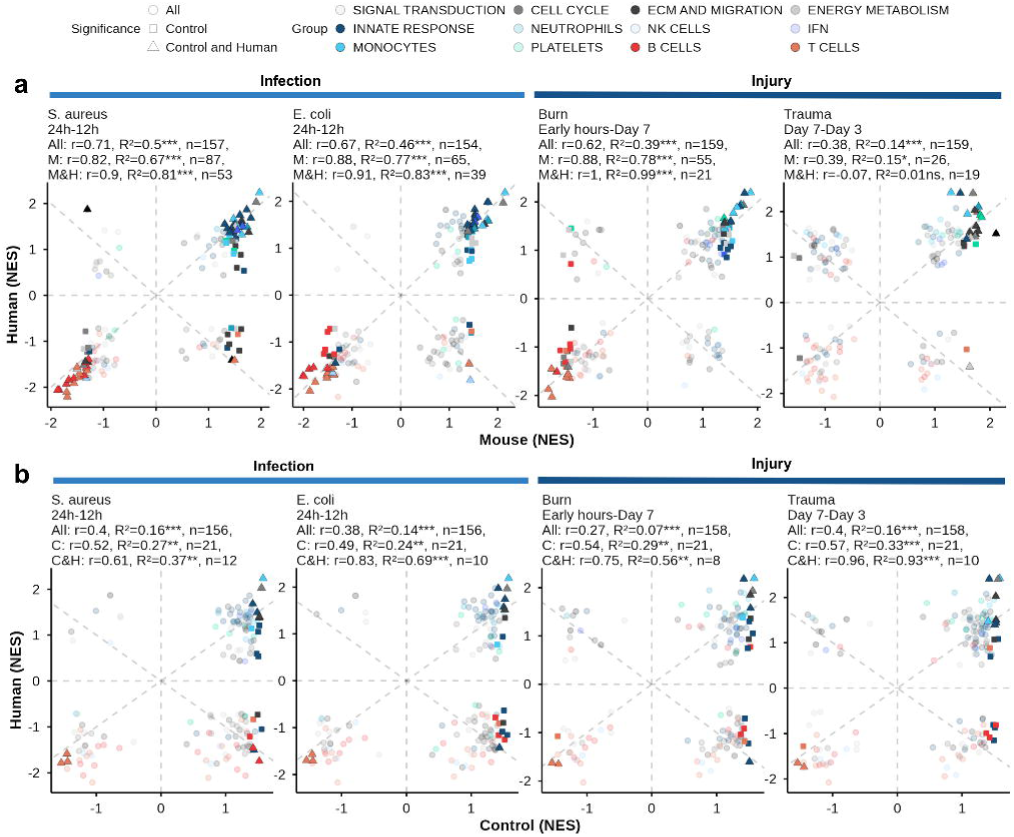
Immune response analysis using the Normalized Enrichment Scores (NES) of genes in Blood Transcriptional Modules (BTMs). (a) Correlation of NES between mice and humans across infection and injury. (b) Correlation of NES between human responses and DMD negative control mice. Pearson r, R2 and significance levels are shown for each matched condition, with modules colour-coded by BTM process category and shaped by significance overlap.

**Supplementary Figure S7.**
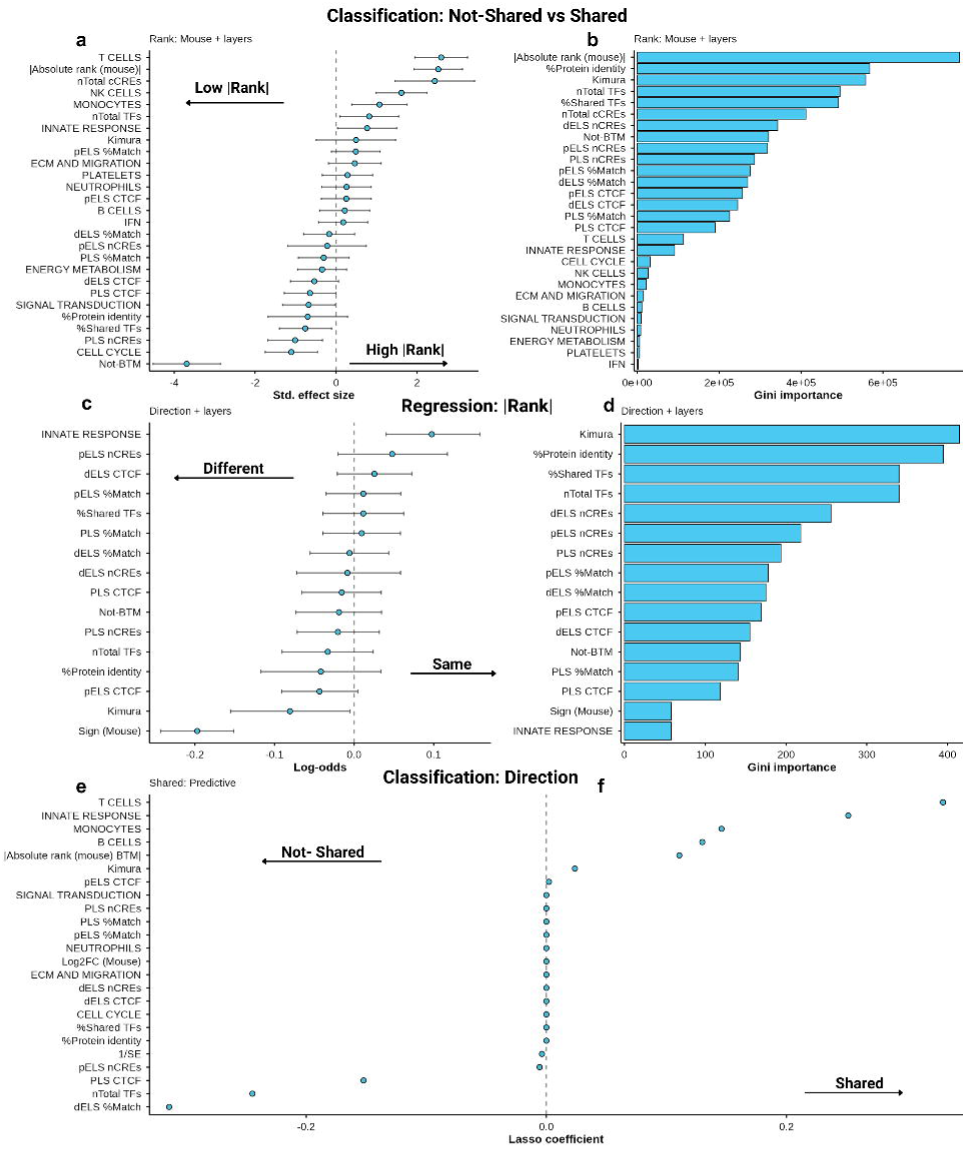
Feature evaluation and importance metrics across multilayered classification and regression models. (a) Standardized effect sizes with 95% confidence intervals from multiple linear regression for human absolute rank. (b) Gini importance from random forest regressors for human rank transfer. (c) Log-odds with 95% confidence intervals from logistic regression for directional concordance. (d) Gini importance from random forest classifiers for directional concordance. (e) Lasso coefficients distinguishing mouse-only from shared LEGs, where positive values favour shared status and negative values favour mouse-only status.

## SUPPLEMENTARY FIGURES

**Supplementary table 1.**
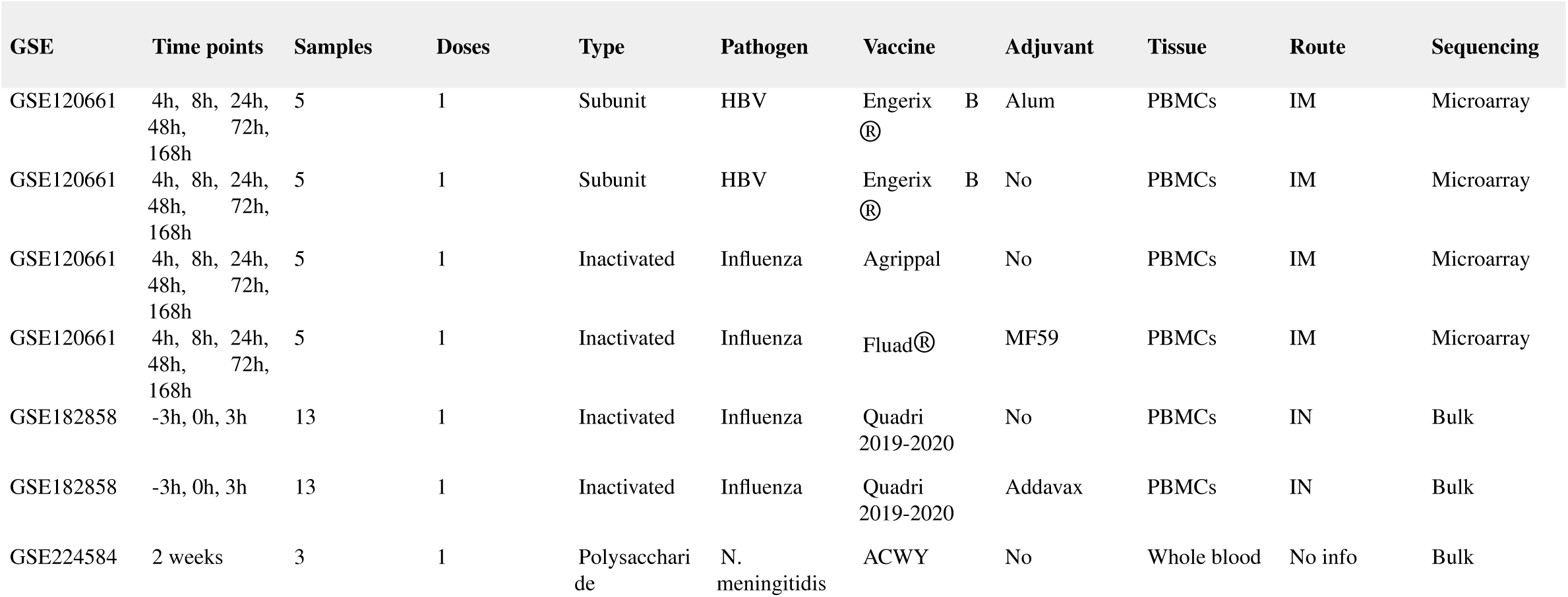
Description of the curated mouse vaccination cohorts. Included studies cover designs ranging from subunit to inactivated vaccines across multiple pathogens (HBV, Influenza, N. meningitidis). Metadata include temporal sampling resolution (hours to weeks), tissue origin (PBMCs, whole blood) and administration routes (IM, intramuscular; IN, intranasal).

---

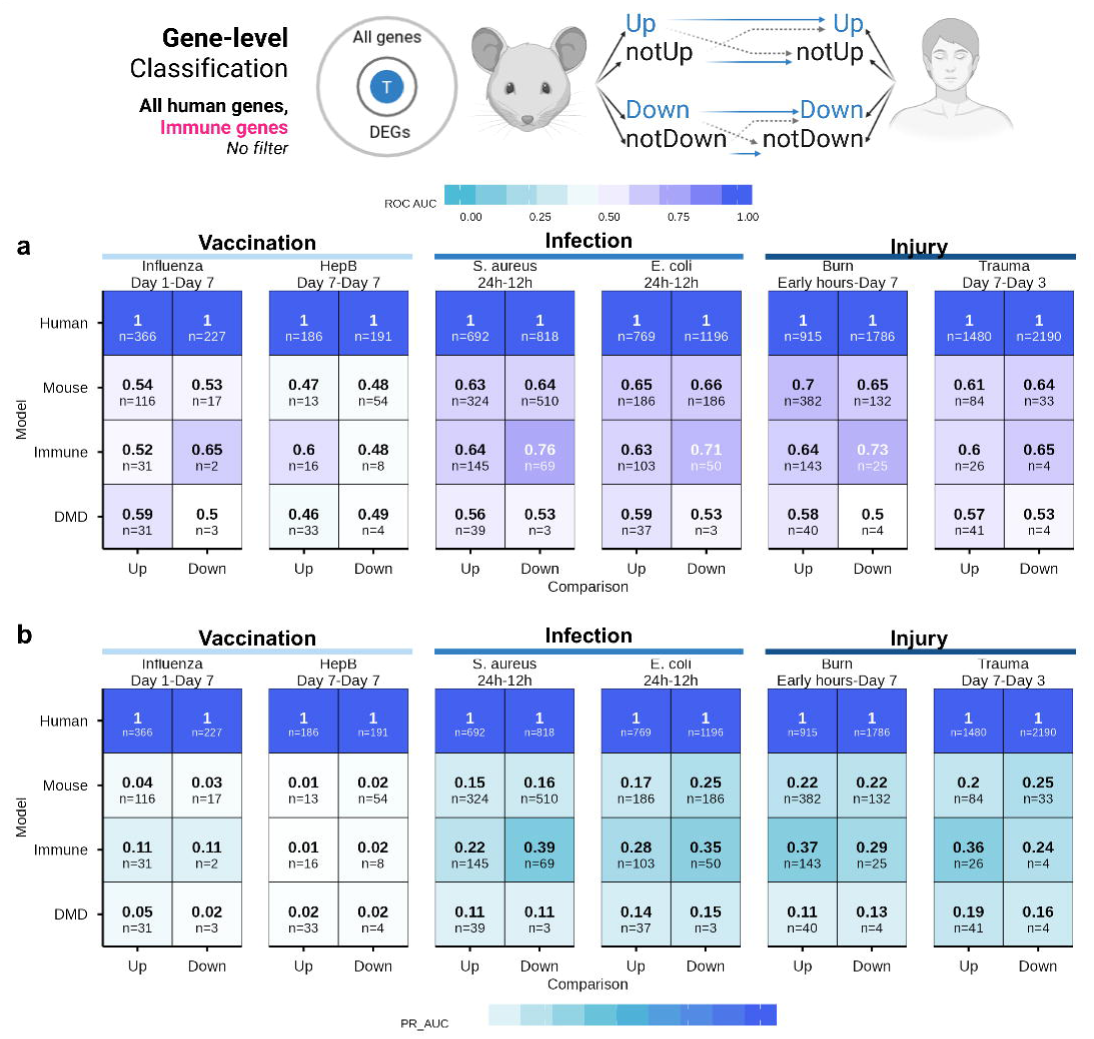

